# Antigen-specific CD8 T cells are generated and reactivated in the bone marrow following viral brain infections and impact the bone marrow niche

**DOI:** 10.64898/2025.12.25.696510

**Authors:** Delaney M. Anani-Wolf, Kelly M. Hotchkiss, Rachael A. Reesman, Christian K. Pfaller, Eliese M. Moelker, Alexandra M. Hoyt-Miggelbrink, Cori E. Fain, Kathryn E. Blethen, Pamela K. Norberg, Shannon E. Wallace, Vatsal Jain, Gerald A. Grant, Mustafa Khasraw, Vidyalakshmi Chandramohan, Aaron J. Johnson, Katayoun Ayasoufi

## Abstract

Bone marrow is the primary immune organ responsible for stem cell maintenance and hematopoiesis. However, the contribution of the bone marrow niche for the generation of adaptive immune responses is less well-understood. We therefore assessed the capacity of virus antigen-specific CD8 T cells to be generated and expanded in bone marrow following acute neurotropic virus infection. Intracranial infection with the neurotropic Theiler’s Murine Encephalomyelitis Virus (TMEV) results in an acute infection in the C57BL/6 mouse which is cleared within 4 weeks post infection due to the generation of an antigen-specific CD8 T cell response against the immunodominant epitope VP2_121-130_. We determined that 1-10% of CD8 T cells present in the femoral, and sternal bone marrow were virus antigen-specific 5-7 days after intracranial TMEV infection. We determined that antigen-specific CD8 T cells were generated in the bone marrow with similar kinetics to that of conventional responses in the secondary lymphoid organs. Importantly, continuous treatment with FTY720, which sequesters T cells outside of the blood, did not eliminate antigen-specific CD8 T cells in the bone marrow compartment indicating generation within the bone marrow niche. Similarly, injection of Poly I:C admixed with fluorescently labeled ovalbumin into the brain generated a similar antigen-specific CD8 T cell response in the bone marrow indicating these responses occur following various brain insults. This model also identified the likely antigen presenting cell (APC) responsible for scavenging antigens from the brain and trafficking into the bone marrow as a migratory myeloid-derived APC. Moreover, following clearance of TMEV infection, CD8 T cells in the bone marrow established durable memory. Antigen-specific memory CD8 T cells in the bone marrow reactivated and expanded upon cognate antigen reencounter. Antigen-specific reactivation of memory CD8 T cells within the bone marrow caused a concurrent increase in Lineage-, Sca-1+, c-Kit+ (LSK) and CD11b+ MHCII+ myeloid cells. This increase in the LSKs and myeloid cells in the bone marrow niche was abrogated with CD8 T cell depletion. We conclude that brain viral infections induce *in situ* effector and memory T cell responses within the bone marrow compartment. Memory recall CD8 T cell responses induce niche dysregulation by expanding LSK cells and lead to an influx of MHCII+ myeloid cells. Our data pave the way for crucial studies of bone marrow resident antigen-specific CD8 T cells in health and diseases.

## Introduction

T cell immunity against CNS malignancies and cancers has historically been perceived to follow conventional T cell priming, infiltration, contraction, and memory formation consistent with peripheral immune responses^1–16^. Brain infections generate antigen-specific T cells that are thought to be primed within the secondary lymphoid organs namely the lymph nodes and the spleen^1–3, 5–13, 16–21^. These antigen-specific T cells infiltrate the brain, interact with brain resident cells, clear the virus, and then contract in numbers. Following contraction, a population of antigen-specific resident memory T cells are generated that remain within the brain parenchyma and exhibit tissue resident memory (TRM) phenotypes^9–11, 15, 17^. Brain TRMs can be reactivated upon cognate antigen reencounter leading to neuroinflammation and neuropathology^11, 13, 15–17, 22–25^. Simultaneously, antigen-specific effector and central memory T cells are also generated that can either circulate through secondary lymphoid organs or enter and survey the brain^17^. We and others have shown that the above concepts largely apply to brain infections following various pathogens including the neurotropic virus Theiler’s murine encephalomyelitis virus (TMEV), Lymphocytic choriomeningitis virus (LCMV), Vesicular stomatitis virus (VSV), and Influenza virus^9, 11, 17, 25^. However, the impact of brain insults on systemic immune function including the primary immune organs is an emerging field. This cross-systems immunological concept is the focus of our study.

Specifically, our group and others have demonstrated that brain insults, including acute infections, TRM reactivation, or brain cancers have systemic consequences making brain insults a systemic disease^17, 26, 27^. Of note, we have shown evidence of systemic lymphopenia, peripheral T cell dysfunction, and spleen and thymic atrophy in the context of brain cancer, acute brain viral infections, sterile injuries, seizures, and more recently following brain TRM reactivation^17, 26–28^. Specific to the bone marrow, we and others have shown that during brain cancer progression T cells enter the bone marrow niche and are sequestrated^26, 27^. To date, it remains unclear how these T cells enter the bone marrow, whether they remain there long-term, or if they impact normal functions of the bone marrow niche. Recently, brain cancers were also shown to impact other systemic functions including bone density and immune responses within several bones including the femur and the skull bones^29^. All in all, brain insults are linked to the dysfunction of peripheral immunity including functions of primary immune organs. However, whether antigen-specific T cells are found within the primary immune organs, namely the bone marrow, following brain viral infections and the extent to which they impact the bone marrow niche acutely and during T cell memory time points remain largely understudied.

The thymus and the bone marrow are primary immune organs in which immune cell development occurs^30–42^. Development of most innate immune cells, B cells, and red blood cells occurs in the bone marrow while the thymus is primarily responsible for T cell development^43, 44^. The bone marrow is exclusively responsible for housing and maintenance of hematopoietic stem and progenitor cells (HSPCs) as well as the process of hematopoiesis^38, 41, 42, 44^. In addition, the bone marrow maintains long-lived plasma cells that secrete antibodies against previously encountered pathogens and vaccinations for the life of the individual^45–48^. As such, bone marrow is typically thought of as a delicate and well-regulated niche with tightly controlled mechanisms to limit entry of various immune cells. Given the inflammatory nature of T cells, it is conceivable that if antigen-specific T cells accumulate in the bone marrow during infections, this may lead to disruptions in hematopoiesis. Given the above, few studies have assessed the capacity for the bone marrow as a site for T cell priming.

In this study, we evaluated the extent antigen-specific CD8 T cells can be generated in the bone marrow compartment during acute neurotropic virus infection. We determined that acute TMEV infection results in early accumulation of antigen-specific CD8 T cells in the bone marrow compartment. Within the bone marrow, antigen-specific CD8 T cells formed durable memory cells weeks after systemic viral clearance. Reencounter with the cognate viral peptide during memory timepoints resulted in renewed expansion and activation of antigen-specific memory CD8 T cells within the bone marrow. Expansion of memory antigen-specific CD8 T cells resulted in proliferation of the HSPCs within the bone marrow signaling possible dysregulation of hematopoiesis. Reactivation of antigen-specific T cells within the bone marrow was also associated with an influx of MHCII+ myeloid cells. Finally, dysregulation of the bone marrow HSPCs plus influx myeloid cells following antigen-specific reactivation of CD8 T cells was abrogated when CD8 T cells were depleted. These findings support the role of the bone marrow resident CD8 T cells as disruptors of hematopoiesis while also governing enhanced influx of MHCII+ myeloid cells into the bone marrow compartment.

## Methods

### Mice and animal care

Male and female C57BL/6 (Cat#000664) mice were purchased from Jackson Laboratory (Bar Harbor, ME). All handling of animals and procedures were approved by the Mayo Clinic and/or Duke University Institutional Animal and Use Committees (IACUC).

### Injection of TMEV

Mice were anesthetized with 2.5% isoflurane and intracranial injection with 10 µL of 2x10^6^ PFU of Daniel’s strain of Theiler’s Murine Encephalomyelitis Virus (TMEV) was given as previously described.^5^ Original TMEV stock was made and provided by Dr. Kevin Pavelko at Mayo Clinic (Rochester, MN) and stored at -80°C.

### Treatment with FTY720

Stock solution of FTY720 (Cat# SML0700-25mg, Millipore/Sigma, Burlington, MA) was diluted in water to inject at a dose of 0.5 mg/kg assuming a 30 g mouse. Randomized, age-matched mice were injected intraperitoneally twice daily with 100 µL of 0.15 mg/mL FTY720 solution beginning two days before TMEV infection and ending on the day of sacrifice. In memory T cell experiments, FTY720 was initiated 5 days (or according to individual figure legends) prior to euthanasia. This treatment strategy was previously used and found to be effective^17^.

### Treatment with αCD8-Depleting Antibodies

A mixture of two αCD8-depleting antibodies was given at either low dose (αCD8LD) or high dose (αCD8HD). For αCD8LD treatment, 10 µg each of clone YTS169 (Cat# BE0117, BioXcell, Lebanon, NH) and clone 53-6.7 (Cat# BE0004-1, BioXcell) were mixed and diluted in 1X PBS to a concentration of 0.1 mg/mL per clone. Intraperitoneal injection of 200 µL was given for 2 consecutive days to deplete peripheral CD8 T cells prior to reactivation with the VP2_121-130_ peptide. For αCD8HD, 200 µg of each clone (YTS169 and clone 53-6.7) were mixed and diluted in 1X PBS to a concentration of 1 mg/mL per clone. Intraperitoneal injection of 400 µL was given for 3 consecutive days to deplete both peripheral and tissue resident CD8 T cells prior to reactivation with the VP2_121-130_ peptide. As previously described and published, aCD8LD depletes CD8 T cells from the blood and lymphoid organs, while aCD8HD depletes CD8 T cells from all tissues^17, 49^.

### Reactivation of VP2_121-130_-specific T cells

Reactivation of antigen-specific T cells was achieved through tail vein injection of 100 µL of the VP2_121-130_ (FHAGSLLVFM) peptide (Gen-Script, Piscataway, NJ, USA) at a concentration of 1 mg/mL in 1X PBS. E7 (RAHYNIVTF) peptide was injected at the same volume and concentration in the control group. Both VP2_121-130_ and E7 peptides are MHCI (D^b^) restricted^3–5, 19, 50, 51^.

### Labeling of Intravascular CD45^+^ Cells

Mice were intravenously injected (into the tail vein) with 6 µg of FITC αCD45 antibody diluted in 1X PBS (total of 300 µl volume was injected into the tail vein), and a period of 3 minutes was allowed for circulating CD45^+^ cells to be marked. 12 µl of αCD45 antibody (35-0451-U100, Tonbo), was mixed with 288 µl of 1X PBS and injected to achieve 6 µg/mouse concentrations. Mice were then sacrificed according to IACUC guidelines using the isoflurane open drop method and perfused intracardially with 30 mLs of 1X PBS.

### Injection of Poly I:C or Poly I:C+ AF488-labeled OVA into the brain

AF488-OVA (Ovalbumin, Alexa Fluor 488 conjugate, Cat# O34781, Invitrogen, Carlsbad, CA) was diluted in 100 µl of water (20 µg/µl). 10 mg of Poly I:C (NC9180242, Invivogen, San Diego, CA) was diluted in 1 mL of water (10 µg/µl). For the Poly I:C alone group, 40 µL of 10 µg/µL Poly I:C was combined with 20 µL of 1X PBS. For the combination group, 80 µL of Poly I:C was mixed with 40 µL of AF488-OVA. Mice were anesthetized with Ketamine (50-100 mg/kg)/Xylazine (5-10 mg/kg) as approved by our IACUC protocol, hair was shaved, head was cleaned with three alternative swabs of iodine and ethanol, mice were placed in a stereotactic frame (Stoelting, Wood Dale, IL), an incision was made over the head, bregma was visualized, specific locations of interest were identified, and a burr hole was drilled over these positions. 3 µL of the appropriate solutions (described above) were injected over 6 minutes (0.5 µl/min) into two different brain locations in each mouse: lateral ventricle (-0.56 A-P, +1 M-L, -2.3 D-V) and cortex (-0.56 A-P, -2 M-L, -1 D-V) using Hamilton syringes (Cat # 14-824-16, Hamilton, Reno, NV). Each injection was 6 minutes, then 1 min rest, syringe recovery, then another injection, 1 min rest, and final syringe retraction. Injections were performed using automatic pumps connected to the frame (World Precision Instruments, Sarasota, FL). Injection needle was kept in position for 1 minute after injection completion before removal and suture. Mice were given ophthalmic ointment, received analgesia with Carprofen (5-20 mg/kg), given antibiotic ointment on top of sutures, and provided heat until recovery. 5-7 days after injection, organs were harvested and flow cytometry was performed. Mice were perfused with 30 mLs of 1X PBS.

### Organ Processing

Organs were procured after iv CD45 labeling, euthanasia, and following intracardial perfusion with 30 mLs of 1X PBS. Euthanasia was performed using isoflurane open drop method as approved by IACUC. After processing as below, cells from all organs were counted by hemocytometer using trypan blue exclusion to identify and exclude dead cells before staining for flow cytometry. To calculate cell counts, the hemocytometer counts were multiplied by the frequency of various population of live cells after gating in Flowjo. For initial processing, samples were filtered through 100 µl filters (130-110-917, Miltenyi Biotec, Bergisch Gladbach, Germany). Samples were filtrated again into flow tubes with restrainer caps before staining (352235, Falcon).

### ACK lysis buffer

8.3 g ammonium chloride, 1 g potassium bicarbonate, and 250 µL of 0.5 M EDTA solution was dissolved in a total volume of 1 L of water (pH to 7.2-7.4). This buffer was made in house and used as ACK lysis buffer.

### Heparin solution and blood collection and processing

Stock heparin solution was made by placing 240 mg of 180 USP/mg heparin powder (H3393-50KU, CAS# 9041-08-1, Millipore/Sigma) in 43.2 mL of water. This creates a 1000 USP/mL stock solution. From the stock solution, a 1:4 dilution with 1X PBS was performed (10 mL stock heparin + 30 mL of 1X PBS) to create a working solution. 1 mL of working solution per 100 µl of blood was used. Blood was collected in 100 µL aliquots in 5-mL flow tubes (352052, Falcon) containing 1 mL of heparin working solution. Blood was processed as previously described^27^. Briefly, the blood heparin mixture was spun at 400 × g for 5 mins, ACK lysed (2 mLs, 3 mins with gentle agitation), quenched (with 1X PBS), spun again (at 400 × g, 5 min), washed one more time (1X PBS), spun again (400 × g, 5 min), and resuspended in 500 µl of 1X PBS for cell count. All spins were performed at 4°C.

### Brain processing

Brains were mechanically homogenized in RPMI by manual glass dounce (Cat # 7727-7, PYREX, Corning, Corning, NY) and myelin was removed with 30% Percoll (P1644-1L, Sigma) gradient as previously described^2, 6, 17, 27, 52^. After myelin aspiration, samples were resuspended into 50 mL total volume of RPMI and spun at 400 × g for 10 mins at 4°C and then aspirated to ∼1 mL volume. Samples were transferred to 1.5-mL microcentrifuge tubes and previous conical tube was rinsed with ∼500 µL of RPMI which was then added to samples in microcentrifuge tubes to ensure maximum cells recovery. Samples were then spun at 1.8 RCF for 10 mins at 4°C, aspirated to the pellet, and resuspended into 500 µL total volume of RPMI for cellular counting. If samples appeared bloody, they were resuspended in 100 µL of ACK lysis buffer as described above for 1 minute. Samples were then quenched with RPMI and centrifuged prior to resuspension in 500 µL RPMI.

### Spleen processing

Spleens were weighed and then mechanically homogenized in RPMI using the frosted sides of two microscopy slides. Samples were spun down (400 × g, 10 mins) and aspirated to the pellet, and red blood cells were lysed using ACK lysis buffer (∼600 µl-1 mL for 1 minute per sample). Buffer was quenched with RPMI, samples were spun again (400 × g, 5 mins) and resuspended in 5 mLs of RPMI for cellular counting.

### Thymus processing

Thymi were weighed and then mechanically homogenized in RPMI using the frosted sides of two microscopy slides. Samples were then spun at 400 × g for 10 mins at 4°C and resuspended in 5 mLs of RPMI for cellular counting.

### Lymph node processing

Lymph nodes were mechanically homogenized in RPMI using the frosted sides of two microscopy slides, then spun at 400 × g for 10 mins at 4°C, and resuspended in 5 mLs of RPMI for counting.

### Femoral Bone Marrow processing

Femoral bone marrow was harvested by cutting off both femoral diaphysis and flushing the medullary space with 1X PBS using a 21-gauge needle attached to a 3-mL syringe. Samples were spun down (400 × g, 10mins) and aspirated to the pellet, and red blood cells were lysed using ACK lysis buffer (∼600 µl-1ml per sample for 1 minute). Samples were quenched with RPMI and spun at 400 × g for 5 minutes at 4°C and resuspended in 5 mLs of RPMI for cellular counting.

### Sternal and Cranial Bone Marrow

Sterna and calvaria were dissected and placed in RPMI. Each sample was cut into strips, and a 21-gauge and a 25-gauge needle with a 3-mL syringe were used to flush bone marrow into a petri dish with 1X PBS. Bones were placed into 0.5-mL microcentrifuge tubes with a hole punched in the bottom and that tube was placed into a 1.5-mL microcentrifuge tube. These were spun at 4 RCF for 2 minutes to help remove marrow from the bone and collect it in the 1.5-mL tube. The pellet was collected, and the bone was flushed for a second time with 1X PBS to remove the remaining marrow. All 1X PBS containing marrow and marrow removed by centrifugation from each sample were combined into a 15-mL conical tube and spun at 400 × g for 10 minutes at 4°C. Samples were aspirated to the pellet and red blood cells were lysed using 100 µL of ACK lysis buffer for 1 minute and quenched with RPMI. They were then spun at 400 × g for 5 minutes at 4°C, aspirated to the pellet, and resuspended in 2 mLs for sternal bone marrow samples and 1 mL for cranial bone marrow samples for cellular counting.

### Flow cytometry

**See Supplementary Table 1** for all antibodies, concentrations, catalog numbers, and unique reagents. Samples were placed in the dark for all staining incubation periods, and all washes were done with 1X PBS. Virus antigen-specific cells were stained with H-2D^b^: VP2_121-130_ (FHAGSLLVFM) tetramer (NIH Tetramer Core Facility, Emory University in Atlanta, GA.) at a concentration of 1:100 for 30 mins at RT and washed once. When K^b:^SIINFEKL tetramer was used, 1:50 concentrations (at RT) were used. Viability staining was done with Zombie UV dye at a concentration of 1:1000 in 1X PBS for 20 mins at RT and then washed once. Master mixes included antibodies in concentrations ranging from 1:50-1:1000 and FC block at a concentration of 1:100 to prevent non-specific binding, brilliant stain buffer plus to prevent brilliant violet dye interactions (Cat # 566385, BD Biosciences, Franklin Lakes, NJ), and monocyte block to prevent antibody phagocytosis by macrophages (True-Stain Monocyte Blocker™, Cat # 426103, Biolegend, San Diego, CA). Master mix staining was done at 4°C for 30 mins and samples were then washed two to three times. Cells were fixed using 250 µL/sample CytoFix Buffer (Cat# 554655, BD Biosciences) at 4°C for 20 mins, then washed once with 1X PBS. Cells were placed in 200 µL of PBS and stored in the dark at 4°C until data collection (maximum storage time of one week) with a 5-laser Cytek Aurora flow cytometer running SpectroFlo® software (Cytek, Fremont, CA).

### Data Analysis and Graphing

Flow cytometry data was analyzed using FlowJo version 10.10.0 (FlowJo LLC, Ashland, OR). Statistical analysis and graphing were done using GraphPad Prism version 10.5.0 (La Jolla, CA). Details of statistical analysis are mentioned in individual figure legends. Data is represented as individual data points with mean ± standard deviation (SD). Statistical significance was determined using Welch’s t-test or ANOVA with post hoc or Mann-Whitney or Kruskal-Wallis with post-hoc analysis as mentioned in individual figure legends. Normality was assessed by the Shapiro Wilk’s test. Outlier analysis was performed using the ROUT test. Experimental design and schema were created using Biorender.com (Created in https://BioRender.com).

## Results

### Virus antigen-specific CD8 T cells expand within the secondary lymphoid organs (SLO) prior to brain infiltration during acute TMEV infection

We sought to evaluate the extent to which antigen-specific CD8 T cells are found in primary immune organs following brain viral infections. C57BL/6 mice were intracranially infected with TMEV. TMEV is a neurotropic virus that causes an acute infection in C57BL/6 mice, leading to the generation of a robust CD8 T cell response against the immunodominant viral antigen, VP2_121-130_, presented in the context of the D^b^ class I molecule^5^. Virus is then effectively cleared from the host 10-21 days post infection beyond which no evidence of viral infectivity is detectible in this strain of mice^2, 5^.

We first evaluated frequencies and numbers of antigen-specific CD8 T cells in the brain, secondary lymphoid tissues (spleen, cervical and inguinal lymph nodes), and primary lymphoid organs (thymus, femoral bone marrow, and sternal bone marrow) on days 1, 2, 3, 4, 5, and 7 post TMEV infection **(Fig 1A)**. To evaluate numbers of D^b^: VP2_121-130_ epitope specific CD8 T cells, we used D^b^: VP2_121-130_ tetramers. As such, D^b^: VP2_121-130_ antigen-specific CD8 T cells against the VP2 antigen are referred to as D^b^: VP2^+^. As expected, at the site of infection, intracerebral infection with TMEV results in the generation of D^b^: VP2^+^ CD8 T cells by day 5 post infection **(Fig 1B)**. In the brain, antigen-specific CD8 T cells are detectible by day 5 and peak on day 7 post infection **(Fig 1B, Fig S1A)**. We previously determined that day 7 is the peak of the D^b^: VP2^+^ CD8 T cell response **(Fig S1A and** ^2, 5, 17, 27, 50, 51, 53^). Classically, antigen-specific CD8 T cells are primed in secondary lymphoid organs (SLO) days prior to the peak infiltration at the site of infection. We hence evaluated frequencies and numbers of D^b^: VP2^+^ CD8 T cells in the draining lymph nodes (cervical lymph nodes, CLN), spleen, and non-draining lymph nodes (inguinal lymph nodes, ILN) on days 1, 2, 3, 4, 5, and 7 post infection in mice intracerebrally infected with TMEV. In the draining lymph nodes, we found minimal numbers of D^b^: VP2^+^ CD8 T cells in the CLN as early as days 1-2 and expanded numbers by days 4-5 **(Fig 1C)**. In the spleen, we found D^b^: VP2^+^ CD8 T cells as early as 3 days post-infection with expanded numbers reaching the highest numbers 7 days post-infection **(Fig 1C, and Fig S1B)**. In the non-draining inguinal lymph node, ILN, D^b^: VP2^+^ CD8 T cells were detectible on days 5-7 post infection **(Fig S1C)**.

**Figure 1:**
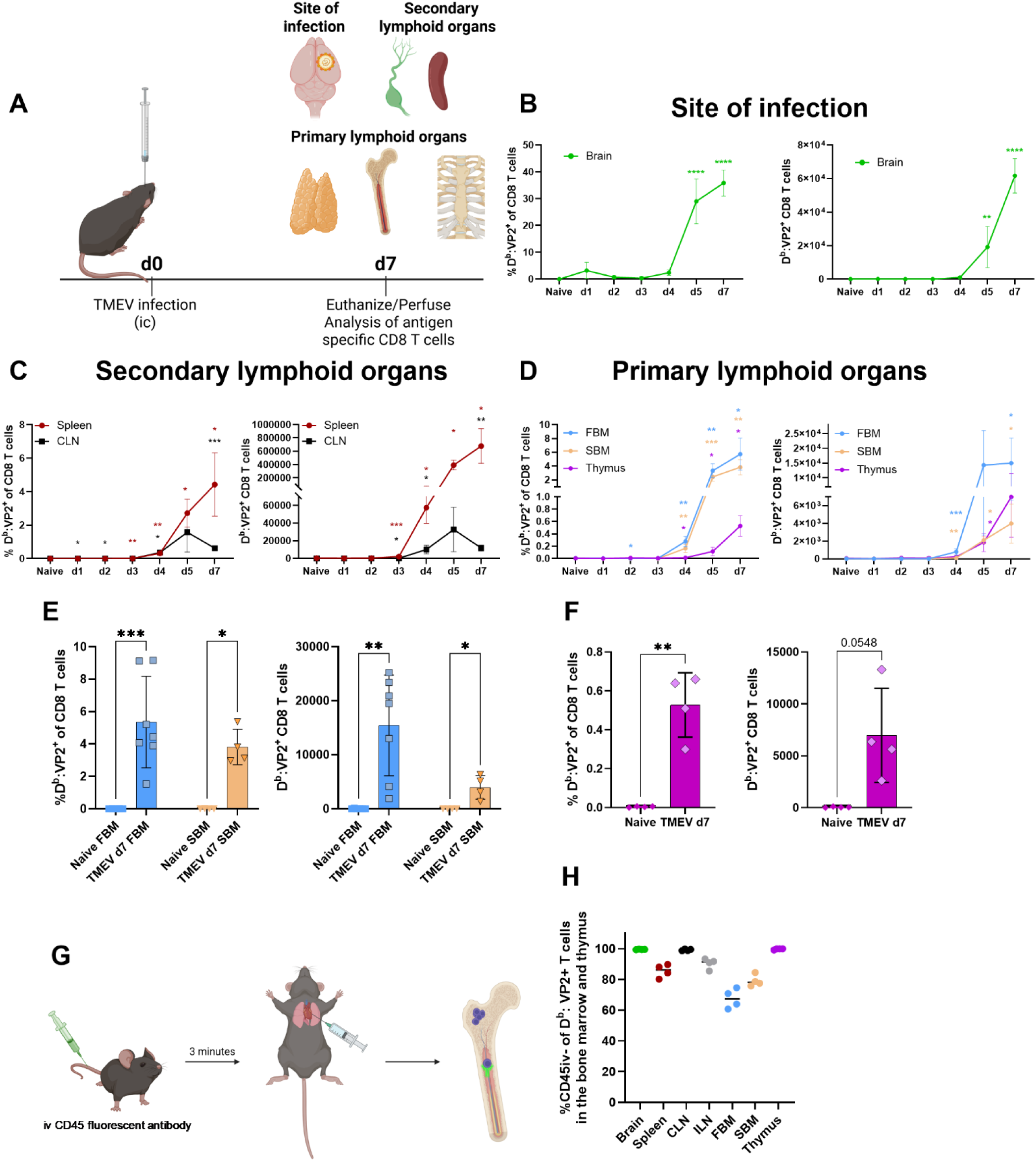
Virus antigen-specific CD8 T cells accumulate in the bone marrow following neurotropic TMEV infection. **A)** Experimental design is shown. C57BL/6 mice were intracerebrally infected with TMEV on day 0. Mice were euthanized and perfused at times between 1-7 days post-infection. Uninfected naïve mice were used as controls. The site of infection (brain), secondary lymphoid organs including draining lymph nodes (cervical lymph nodes, CLN), non-draining lymph nodes (inguinal lymph nodes, ILN), and spleen and primary immune organs including the thymus, and two sources of the bone marrow (femoral, and sternal) were collected, processed, and analyzed via spectral flow cytometry. **B)** Kinetics of the appearance of D^b^: VP2+ CD8 T cells in the brain is shown by frequencies (of CD8 T cells) and absolute counts. Antigen-specificity is measured via tetramer staining. In the brain, antigen-specific CD8 T cell response peaks on day 7 post TMEV infection. **C)** Kinetics of the appearance of D^b^: VP2+ CD8 T cells in the SLO is shown. Frequencies and numbers of antigen-specific CD8 T cells are quantified. As expected, D^b^: VP2+ CD8 T cells appeared and expanded on days between 1-7 post infection in the SLO. **D)** Kinetics of the appearance of D^b^: VP2+ CD8 T cells in the bone marrow and thymus are shown. Frequencies and numbers of antigen-specific CD8 T cells are quantified. Surprisingly, D^b^: VP2+ CD8 T cells are also found and expanded in primary immune organs on days between 2-7 post-infection. **E-F)** Frequencies and numbers of D^b^: VP2+ CD8 T cells in the bone marrow **(E)**, and the thymus **(F)** are shown 7 days post TMEV infection. **G)** Mice were intravenously injected with a fluorescently labeled αCD45 antibody (CD45iv). Three minutes later, mice were euthanized and perfused with 30mLs of PBS. Organs were then harvested, and D^b^: VP2+ CD8 T cells were analyzed for the expression of the iv-label. **H)** The majority of the D^b^: VP2+ CD8 T cells in the brain, SLO, and the primary immune organs are iv- meaning that they are located within the tissue parenchyma (and not in/near vasculature). In **B-D**, a Two-Way ANOVA (or Mixed model) is performed in which time points are compared. The results were found significant. Post-hoc analysis (Fisher’s LSD test) was then performed to evaluate comparisons between naïve and each time point. In **E**, a one-way ANOVA or a Kruskal-Wallis test (depending on parametric or non-parametric distribution of the data) was performed and found to be significant. Šídák’s or Dunn’s multiple comparisons tests were then performed comparing the pairs shown. Data are shown as individual mice. Mean and SD are shown. In F, an unpaired two-tailed Welch’s t test was performed. ns = P > 0.05, * = P ≤ 0.05, ** = P ≤ 0.01, *** = P ≤ 0.001, and **** = P ≤ 0.0001.

### Antigen-specific CD8 T cells are found within the primary immune organs acutely following brain viral infections

Having established that D^b^: VP2^+^ CD8 T cells are generated within 3 days in the SLOs and infiltrate the brain between 5-7 days, we next evaluated if antigen-specific CD8 T cells are also found in the primary immune organs at time points similar to the SLO. We evaluated two sources of the bone marrow (femoral and sternal) and the thymus as the primary immune organs evaluated post TMEV infection. Surprisingly, we found significant numbers of D^b^: VP2^+^ CD8 T cells in the bone marrow between days 2-4, with further expansion occurring between days 4-7 **(Fig 1D)**. D^b^: VP2^+^ CD8 T cells were also detectible in the thymus between days 5-7 **(Fig 1D)**. We next quantified the response in the primary immune organs between uninfected mice and TMEV infected mice on day 7 post infection (peak of the immune response in this infection). In both femoral and sternal bone marrows, significant frequencies and numbers of D^b^: VP2^+^ CD8 T cells were found in infected mice compared to the uninfected controls **(Fig 1E)**. Similarly, significant frequencies of D^b^: VP2^+^ CD8 T cells were found in the thymus 7 days post-infection compared to the uninfected controls **(Fig 1F)**. Numbers of D^b^: VP2^+^ CD8 T cells in the thymus did not reach a statistically significant difference when compared to the uninfected controls (p=0.0548, **Fig 1F**). We next evaluated whether the D^b^: VP2^+^ CD8 T cells in the brain, SLO, and the primary immune organs were in or near vasculatures or found within the tissue parenchyma using iv labeling with intravenously injected fluorescently labeled αCD45 antibody into mice 3 minutes before euthanasia. Immune cells inside tissues or away from the vasculature will not be labeled with intravenously administered αCD45 antibody and will be referred to as iv- **(Fig 1 G)**. Mice were injected with the iv-label, euthanized, and then perfused prior to organ harvest **(Fig 1 G)**. In the brain, and SLO, over 80-100% of D^b^: VP2^+^ CD8 T cells were iv- indicating the majority of antigen-specific CD8 T cells are tissue resident **(Fig 1H)**. In the bone marrow, we found that over 50% of the D^b^: VP2^+^ CD8 T cells were iv- while nearly 100% of D^b^: VP2^+^ CD8 T cells in the thymus were iv- **(Fig 1H)**. Therefore, virus antigen-specific CD8 T cells are found within the primary immune organs following acute neurotropic TMEV infection and expand with similar kinetics to those of the SLO. In addition, the majority of the D^b^: VP2^+^ CD8 T cells in the brain, SLO, and the primary immune organs are tissue resident.

### Virus antigen-specific CD8 T cells are generated within the bone marrow niche following neurotropic TMEV infection

Thus far we have observed increased levels of virus antigen-specific CD8 T cells accumulating within the bone marrow niche during acute TMEV infection. We next tested if these antigen-specific CD8 T cells are generated within the bone marrow *in situ*, or if they are generated in the SLO and infiltrate the marrow subsequently. To answer this question, we used FTY720, which is a known sphingosine 1 phosphate (S1P) modulator and sequestrates T cells within the SLO effectively reducing trafficking through the blood^54–56^. FTY720 was initiated two days before TMEV infection. FTY720 was continuously given twice a day for the entire duration of the experiment. Vehicle (water) treated animals were used as control. Experimental groups were then euthanized on day 7 post-infection **(Fig 2A)**. As expected, FTY720 treatment reduced frequencies and counts of total T cells, CD4, and CD8 T cells in the blood indicating significantly inhibited T cell trafficking **(Fig S2A-C)**. Having verified that FTY720 works as expected in the blood and sequesters T cells within the SLO, we next evaluated frequencies and numbers of D^b^: VP2^+^ CD8 T cells in the draining and non-draining lymph nodes (CLN and ILN) and the spleen respectively. We found that antigen-specific CD8 T cells were found in both infected groups regardless of FTY720 treatment. While both infected groups clearly had detectible D^b^: VP2^+^ CD8 T cells, the difference to corresponding naïve groups did not reach statistical significance in the lymph nodes. Regardless, FTY720 treated mice demonstrated significantly increased frequencies and numbers of D^b^: VP2^+^ CD8 T cells on day 7 in all lymph nodes indicating T cell sequestration and lack of trafficking out of the SLO **(Fig 2B-C)**. In the spleen, we found significant frequencies of D^b^: VP2^+^ CD8 T cells in both infected groups regardless of FTY20 treatment **(Fig 2D)**. Interestingly, FTY720 significantly reduced D^b^: VP2^+^ CD8 T cells in the spleen of TMEV infected mice indicating that the majority of spleen D^b^: VP2^+^ CD8 T cells are being contributed by lymph nodes through the blood circulation **(Fig 2D)**. This is corroborated by the data presented in **Fig 2B-C** in which FTY720 treatment results in increased accumulation of D^b^: VP2^+^ CD8 T cells in the lymph nodes.

**Figure 2:**
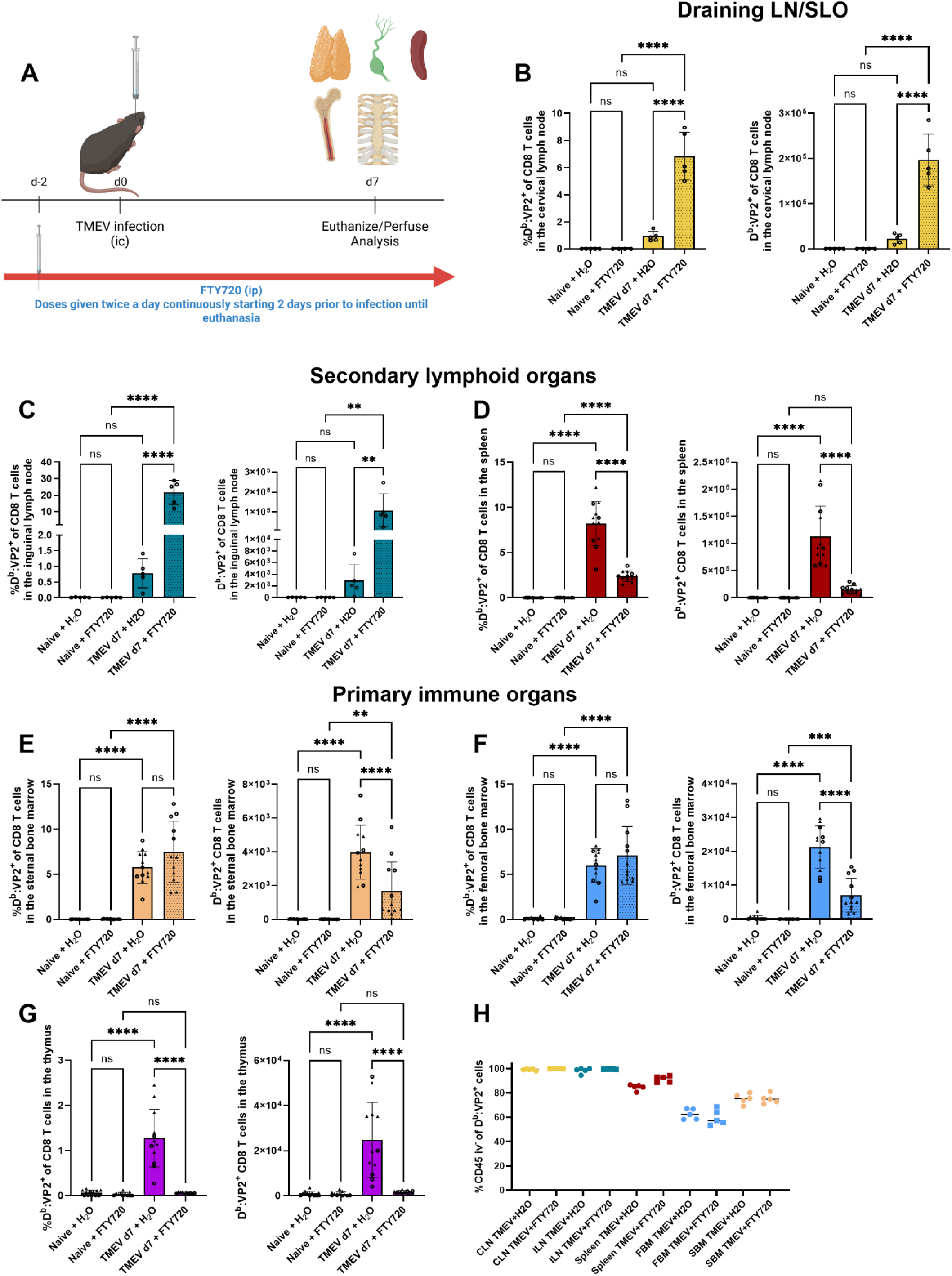
Virus antigen-specific CD8 T cells are generated in the bone marrow within seven days of acute TMEV infection. **A)** Experimental design is shown. Mice were infected with TMEV in the brain and organs were harvested 7 days later. FTY720 was given intraperitoneally starting 2 days before the infection. FTY720 treatment was done twice a day (continuously). Control mice received vehicle (water). **B)** Frequencies and numbers of D^b^: VP2+ CD8 T cells in the cervical lymph nodes are quantified via tetramer staining and flow cytometry. **C)** Similarly, frequencies and numbers of D^b^: VP2+ CD8 T cells in the inguinal lymph nodes are quantified. **D)** Frequencies and numbers of D^b^: VP2+ CD8 T cells in the spleen are quantified. **E)** Frequencies and numbers of D^b^: VP2+ CD8 T cells in the femoral bone marrow are quantified. **F)** Frequencies and numbers of D^b^: VP2+ CD8 T cells in the sternal bone marrow are quantified. **G)** Frequencies and numbers of D^b^: VP2+ CD8 T cells in the thymus are quantified. **H)** Mice received an intravenous injection of a fluorescently labeled αCD45 antibody 3-minutes prior to euthanasia and perfusion. Frequencies of iv- D^b^: VP2+ CD8 T cells are quantified. Cells negative for the iv-label were quantified within the D^b^: VP2+ CD8 T cell gate. In **B-G**, a one-way ANOVA followed by Holm-Šídák’s multiple comparisons test was performed comparing the designated pairs. Data are shown as individual mice. Mean and SD are shown. ns = P > 0.05, * = P ≤ 0.05, ** = P ≤ 0.01, *** = P ≤ 0.001, and **** = P ≤ 0.0001.

Having established that FTY720 sequesters T cells away from the blood, we next asked if D^b^: VP2^+^ CD8 T populations in the bone marrow were impacted. Interestingly, frequencies of D^b^: VP2^+^ CD8 T cells in the femoral or sternal bone marrows were similar in TMEV infected groups regardless of treatment with FTY720 **(Fig 2E-F)**. Numbers of D^b^: VP2^+^ CD8 T cells in the bone marrow were marginally reduced following FTY720 treatment **(Fig 2E-F)**. Regardless, similar frequencies of D^b^: VP2^+^ CD8 T cells were generated in the bone marrow at a rate similar to the SLO but independent of infiltration from circulation. Given that frequencies of D^b^: VP2^+^ CD8 T cells were similar in all sources of the bone marrow in FTY720 treated TMEV infected mice, but reduced in the spleen implies the spleen contributes to circulation of D^b^: VP2^+^ CD8 T cells while bone marrow does not. This data also indicates that D^b^: VP2^+^ CD8 T cells are generated in the bone marrow without recruitment from circulation.

We also evaluated the generation of D^b^: VP2^+^ CD8 T cells in the thymus, another primary immune organ. We found that FTY720 treatment completely abrogated D^b^: VP2^+^ CD8 T cells in the thymus indicating that these cells infiltrate the thymus from circulation and are not primed *in situ* **(Fig 2G)**. Finally, we evaluated whether D^b^: VP2^+^ CD8 T cells were located in or near vasculature using iv-labeling as previously described in **Fig 1G**. We found that the majority of D^b^: VP2^+^ CD8 T cells were located away from the vasculature and were iv- and this was not impacted by FTY720 **(Fig 2H)**. In summary, we determined that antigen-specific CD8 T cells are generated *in situ* within the bone marrow niche following brain viral infections and not transported from the SLO.

### Antigen introduced into the brain is preferentially scavenged by myeloid APCs

Thus far, acute neurotropic viral infections in the brain resulted in the accumulation of viral antigen-specific CD8 T cells within the bone marrow niche. We next evaluated if this phenomenon was specific to viral infections or more ubiquitous to brain insults in general. To avoid caveats of neuronal damage from an actual viral infection, we intracranially administered AF488-labeled-OVA peptide antigen with Poly I:C (OVA-PIC ic) into the brain. Poly I:C was administered alone into the brain as a negative control **(Fig 3A)**. First, we sought to establish the state of neuroinflammation in the brain and evaluate if OVA-PIC ic generates antigen-specific CD8 T cells within the brain using H2-K^b^: SIINFEKL tetramer staining to quantify OVA-specific CD8 T cells.

**Figure 3:**
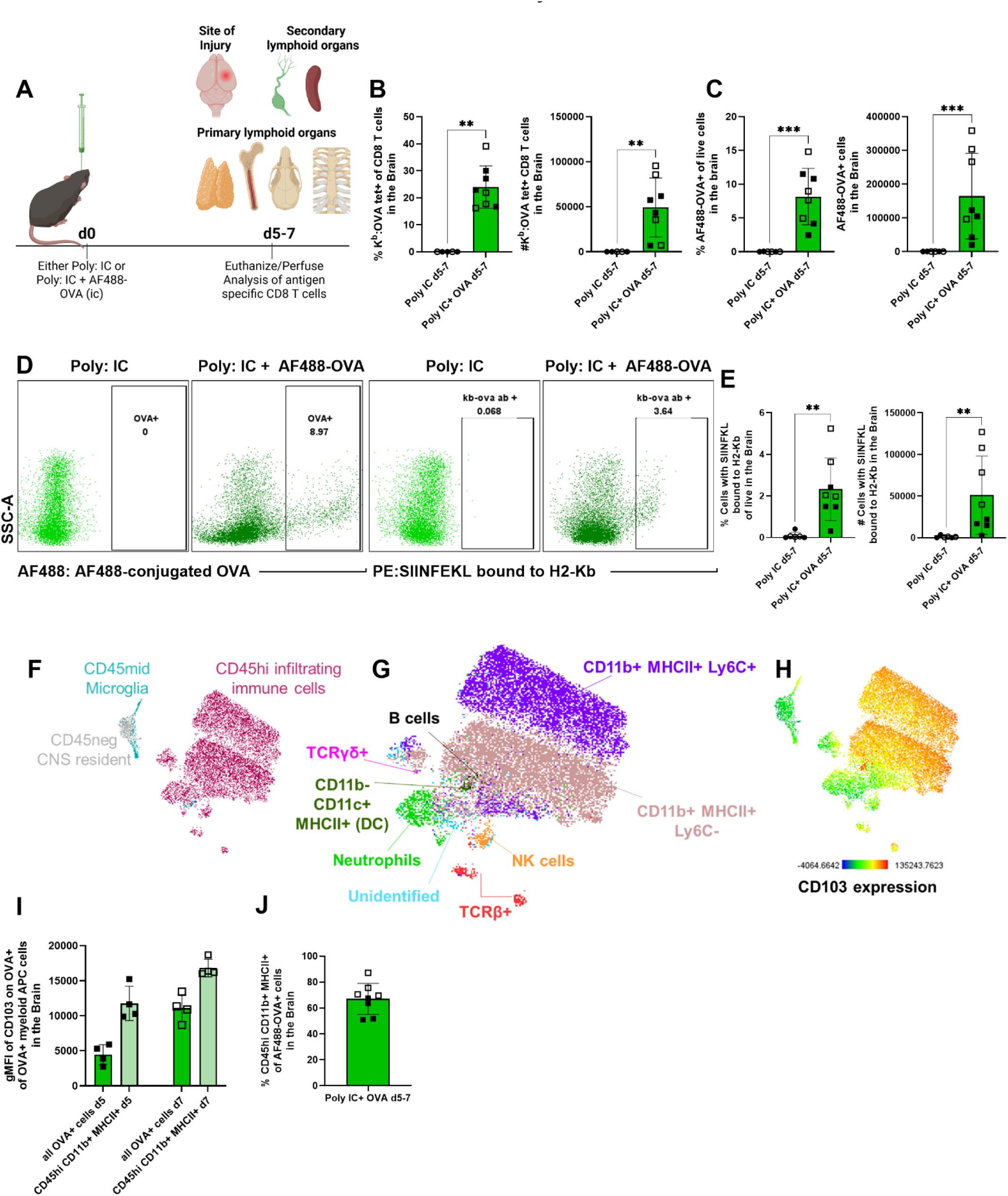
Infiltrating myeloid APCs scavenge antigens within the brain during acute intracranial inflammatory insults. **A)** Experimental design is shown. **B)** OVA specific CD8 T cells within the brain are quantified. **C)** Frequencies and numbers of cells with detectible OVA in them are quantified. **D)** Representative flow plot shows AF488+ cells within the brain on the left. On the right, cells staining positive for the CD8 OVA epitope within MHCI are quantified. **E)** Frequencies and numbers of SIINFEKL-H2-K^b^ (cells presenting the CD8 OVA epitope within MHCI) in the brain are quantified. **F)** Total AF488+ (cells that have OVA in them) cells were concatenated, and a UMAP was built. Majority of OVA+ cells are CD45^hi^ infiltrating immune cells. **G)** Further analysis of cell types indicates that the majority of OVA+ CD45^hi^ cells are APCs especially those originating from myeloid cells (CD11b+). **H)** CD103 expression is shown indicating DCs and myeloid APCs that are OVA+ have the highest expression indicating migratory potential. **I)** Geometric mean fluorescent intensity of CD103 expression is shown in all AF488+ cells and in myeloid APCs. **J)** Over 50% of AF488+ cells were CD11b+ MHCII+ CD45^hi^ cells consistent with myeloid APC identity. Open squares/circles are data from day 7, while closed squares/circles are data from day 5 post-injury. Data are shown as individual mice. Mean and SD are shown. For comparisons in **B**, **C**, and **E**, a Mann Whitney test was used comparing the groups. Comparisons made are shown on the graph. ns = P > 0.05, * = P ≤ 0.05, ** = P ≤ 0.01, *** = P ≤ 0.001, and **** = P ≤ 0.0001.

We determined that OVA-PIC i.c generated significant numbers of K^b^:OVA specific CD8 T cells within the brain 5-7 days post intracranial injection **(Fig 3B)**. We next tested the presence of OVA protein, in the brain at this timepoint by using AF488 as a surrogate marker. Flow cytometry analysis showed AF488 expression in a significant number of CD45+ cells indicating that OVA has been phagocytosed and retained within the brain 5-7 days post injection **(Fig 3C-D)**. In addition, we evaluated the presence of the processed SIINFEKL peptide within MHCI using H2-K^b^-bound-to-SIINFEKL antibodies as this reagent detects the OVA epitope presented to CD8 T cells within the MHCI H2-K^b^. We detected significant numbers of cells that stain positive for SIINFEKL-H2-K^b^ antibody, indicating that these cells picked up, processed and presented OVA in the context of the K^b^ MHC class I molecule **(Fig 3D-E)**. We next defined the identity of the cells that were AF488+ (OVA+). We found that the overwhelming majority of the OVA+ cells were within CD45^hi^ infiltrating immune cells. Only a minority of cells were consistent with being microglia and CD45 negative CNS resident cells **(Fig 3F)**. The majority of the cells that were OVA+ were also CD11b+ and MHCII+ indicating their identity to be of myeloid origin **(Fig 3G)**. We next evaluated CD103 expression, a marker associated with migratory potential, on AF488+ cells within the brain and found that CD11b+ MHCII+ myeloid APCs and CD11b-CD11c+ MHCII+ DCs expressed high levels **(Fig 3H**, MFI quantified **in 3I)**. Finally, our quantification revealed that over half of OVA+ cells in the brain were of myeloid origin and MHCII+ indicating APC potential (descriptive statistics for % CD11b+ MHCII+ CD45^hi^ of AF488+ includes minimum 51, maximum 87.2, mean 67.09, SD 11.97) **(Fig 3J)**. All in all, we have successfully developed a model in which antigen-specific responses within the brain and antigen pick up are recapitulated without the need for a live virus and/or the associated damage from an actual viral infection. In this model, fluorescently labeled OVA was mixed with Poly I:C and intracranially delivered. At a similar time point to the acute TMEV infection (5-7 days later), we found antigen-specific CD8 T cells within the brain parenchyma. Additionally, we uncovered evidence of phagocytosis and antigen presentation by DCs and myeloid APCs at this time point. Finally, APCs that picked up the antigen from the brain expressed high levels of CD103, indicating migratory capacity.

### Acute non-infectious inflammatory insults delivered within the brain result in the accumulation of antigen-specific CD8 T cells within the bone marrow niche

Thus far, we have shown TMEV infection in the brain results in the generation of antigen-specific CD8 T cells in the bone marrow. We next sought to evaluate if non-infectious insults in the brain also produce a similar antigen-specific T cell responses within the bone marrow. To test this, we again employed our approach to deliver antigen with Poly I:C admixed with AF88-labeled-OVA protein. This novel approach resulted in the generation of antigen-specific CD8 T cells within the brain which was largely associated with antigen being scavenged by myeloid APCs expressing CD103 indicating migratory potential **(Fig 3)**. First, we evaluated the presence of antigen-specific CD8 T cells in the SLO. We found significantly higher frequencies of antigen-specific CD8 T cells in the spleen and CLN 5-7 days post Poly I:C-AF488-labeled OVA delivery into the brain compared to Poly I:C alone groups **(Fig 4A-B)**. Accordingly, we found significant frequencies of AF488+ cells, indicating cells with internalized OVA, in both the spleen and the CLN **(Fig 4C).** These results further validate the model and demonstrate similarities with the TMEV model in terms of the responses within the brain and the SLO.

**Figure 4:**
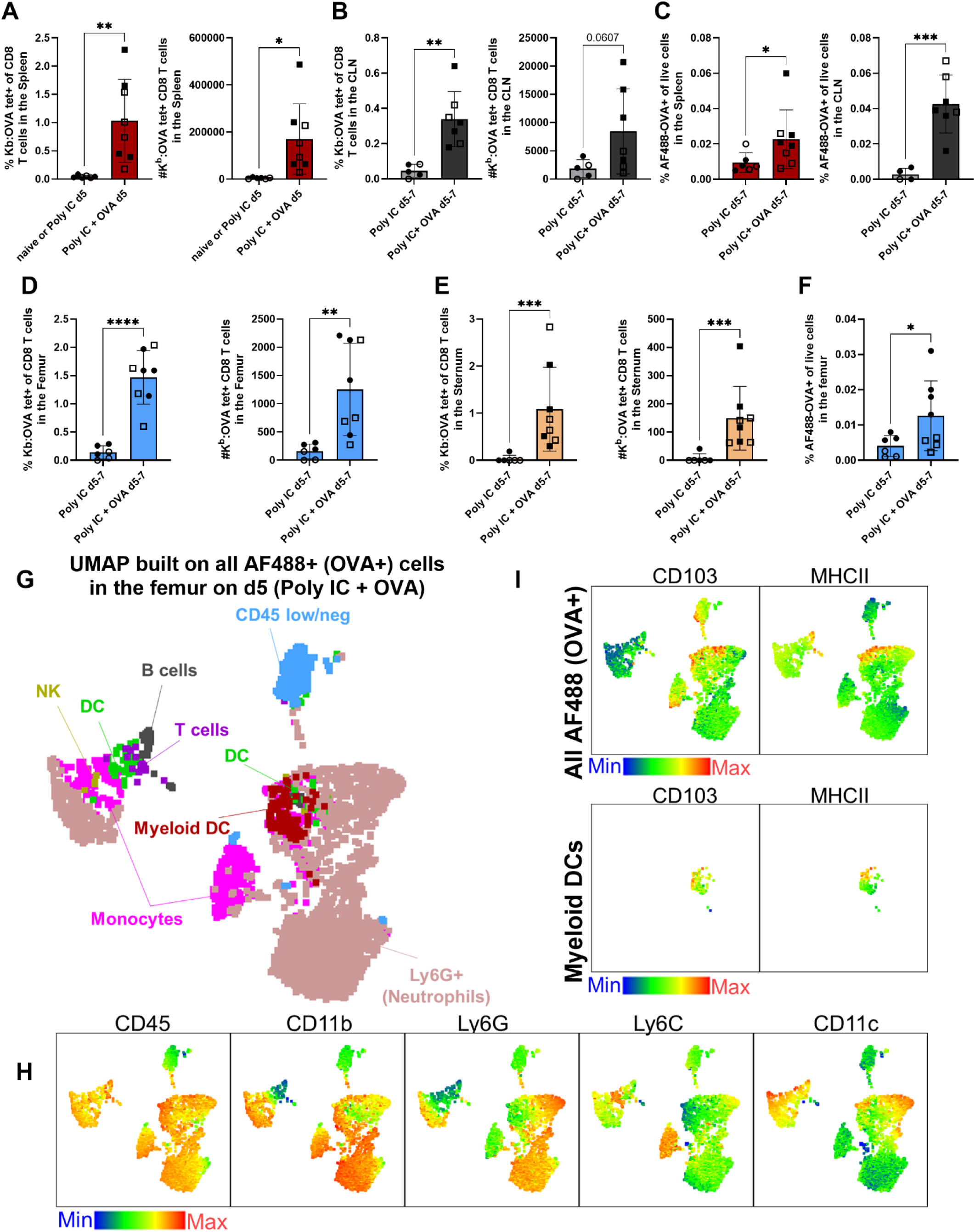
Acute inflammatory brain insults result in the generation of both peripheral and bone marrow derived antigen-specific CD8 T cells. Mice were injected with Poly I:C or Poly I:C+AF488-labeled OVA according to **Fig 3A**. In these mice, the presence of antigen-specific cells and AF488+ cells in the SLO and the bone marrow was evaluated using flow cytometry. **A)** Frequencies and numbers of K^b^:SIINFEKL specific CD8 T cells in the spleen are significantly increased in Poly I:C+OVA group compared to Poly I:C alone groups. **B)** Frequencies and numbers of OVA-specific CD8 T cells in the cervical lymph nodes (CLN) are shown. **C)** Frequencies of AF488+ cells of total lives cells are quantified in the spleen and CLN and found to increase in OVA-injected groups over controls using flow cytometry. **D)** Frequencies and numbers of OVA-specific CD8 T cells in the femoral bone marrow are significantly increased in Poly I:C+OVA group compared to the Poly I:C alone group. **E)** Frequencies and numbers of K^b^: OVA specific CD8 T cells in the sternal bone marrow are significantly increased in Poly I:C+OVA group compared to the Poly I:C alone group. **F)** Frequencies of AF488+ cells of total live cells are quantified in the femoral bone marrow. **G)** AF488+ cells in the femoral bone marrow of mice 5 days following Poly I:C+ AF488-OVA injection were concatenated. UMAP was built on the concatenated AF488+ cells from the femoral bone marrow using FlowJo. Various immune cells were identified using traditional gating strategies. **I)** Expression levels of CD103 and MHCII are shown on all AF488+ cells **(top)**. Expression levels of CD103 and MHCII on AF488+ myeloid DCs are shown **(bottom)**. **H)** Expression levels of CD45, CD11b, Ly6G, Ly6C, and CD11c on all AF488+ cells are shown. All experiments were performed twice (once harvesting organs on day 5 and once on day 7 post-injection into the brain). Data from both experiments are pooled. Open squares/circles are data from day 7, while closed squares/circles are data from day 5 post-injury. Data are shown as individual mice. Mean and SD are shown. For comparisons in **A-F**, either a Welch’s t-test or a Mann Whitney test was used comparing the groups depending on normal or not-normal distribution of data determined by the Shapiro-Wilk’s test. Comparisons made are shown on the graph. ns = P > 0.05, * = P ≤ 0.05, ** = P ≤ 0.01, *** = P ≤ 0.001, and **** = P ≤ 0.0001.

We next evaluated if our approach of non-infectious antigen delivery into the brain also results in the accumulation of antigen-specific CD8 T cells within the bone marrow niche. We identified increased frequencies and numbers of K^b^:SIINFEKL specific CD8 T cells in the femoral and sternal bone marrow of Poly I:C-AF488-OVA injected mice when compared to Poly I:C alone treated animals **(Fig 4D** shows the femoral bone marrow and **4E** shows the sternal bone marrow results). In thymus, we found increased frequencies of K^b^:SIINFEKL specific CD8 T cells (with trending increased numbers) between Poly I:C-AF488-OVA injected mice and Poly I:C groups **(Fig S3A)**. The lack of statistical significance in the thymus K^b^:SIINFEKL specific CD8 T cells cell counts could be due to the thymic involution as a result of brain injury which we have previously published in several models^27^ and also occurred in this model of brain injury **(Fig S3B)**. While we found increased frequencies of AF488+ cells in the femurs compared to controls **(Fig 4F)**, these results did not reach statistical significance in the thymus or sternal bone marrow **(Fig SC-D)**. In the thymus, this is likely due to the fact that cells traffic from the SLO as previously seen in the TMEV model when using FTY720 **(Fig 2G)**. In the sternal marrow, we likely did not find increased frequencies of AF488+ cells due to the general rarity of these cells. Interestingly, we also evaluated the presence of antigen-specific CD8 T cells following OVA delivery in the skull bone marrow. We found significantly increased frequencies and numbers of K^b^: OVA specific CD8 T cells in the skull bone marrow **(Fig S3E)**. Similar numbers of cells were recovered between Poly I:C and Poly I:C+ AF488-OVA groups **(Fig S3F)**. Interestingly, frequencies of AF488+ cells in the skull bone marrow were significantly increased in Poly I:C+ AF488-OVA groups compared to Poly I:C alone controls **(Fig S3G)**. This could be due to the proximity of the skull to the injection site and heavy trafficking of myeloid APCs through brain-skull channels or antigen drainage from the site of injury. Total cells recovered between groups were similar in most organs with the exception of the thymus, where OVA-injected mice had smaller thymi and the spleen where OVA-injected mice had larger spleens compared to Poly I:C alone injected controls **(Fig S3H and S3B)**.

### The majority of antigen-bearing cells bringing brain draining antigens into the femoral bone marrow are myeloid-derived cells in origin including myeloid-derived APCs with migratory potential

Given that antigen-specific CD8 T cells specific for OVA along with AF488+ (OVA+) cells were both detected within femoral bone marrow, we next evaluated the identity of AF488+ cells as the cellular source of antigen. We concatenated all AF488+ cells in the femoral bone marrow of Poly I:C+ AF488-OVA injected mice 5 days post brain injections and built a UMAP on these cells. We determined that the majority of AF488+ cells were of myeloid origin (CD11b+), including neutrophils, monocytes, and myeloid DCs **(Fig 4G)**. Interestingly, few T cells, B cells, NK cells, CD11b-DCs, and CD45 low/neg cells were also amongst cells carrying the OVA antigen within the bone marrow **(Fig 4G)**. We next evaluated the migratory and APC potential of AF488+ cells and the myeloid DC subset by evaluating CD103 and MHCII expression. CD103 was highly expressed on some monocytes, myeloid DCs, and some CD45 low/neg cells **(Fig 4I)**. MHCII was expressed on subsets of DCs, myeloid DCs, and B cells, indicating these cells are likely antigen-presenting cells within the bone marrow **(Fig 4I)**. Finally, comprehensive analysis revealed AF488+ cells are also CD45+, CD11b+, Ly6G+, Ly6C+, and CD11c+, demonstrating that the majority of cells presenting brain introduced OVA antigen are of myeloid origin **(Fig 4H)**.

### Antigen-specific CD8 T cells generated within the bone marrow following neurotropic TMEV infection form durable memory within 60 days

Thus far, we demonstrated that acute neurotropic viral infections and non-infectious acute brain insults result in the accumulation of antigen-specific CD8 T cells within the bone marrow niche. We next sought to determine if these antigen-specific CD8 T cells generate durable memory within the bone marrow. To evaluate memory responses within the bone marrow, we used TMEV infection. Mice were intracranially infected with TMEV and allowed 55 days to clear the virus and form memory T cells. Viral clearance occurs by day 21 post-infection^2, 3, 5, 6, 51, 57^. Groups of mice received FTY720 to reduce T cell trafficking from the SLO into the blood 5 days before euthanasia (given continuously twice a day). Two days before euthanasia, mice received either the E7 peptide (control peptide that does not reactivate TMEV specific CD8 T cells), or the VP2 peptide (virus peptide that activates TMEV specific CD8 T cells) intravenously **(Fig 5A)**. A second group of mice received low dose CD8 (aCD8LD) depleting antibodies two days before peptide reactivation to deplete CD8 T cells. Antigen-specific CD8 T cell responses were evaluated in the femoral bone marrow 2 days after CD8 memory T cell reactivation via systemic VP2 peptide administration. Neither FTY720 treatment nor CD8 depletion impacted total blood cell counts **(Fig S4A)**. CD8 T cell depletion and FTY720 both significantly reduced CD8 T cell frequencies and counts **(Fig S4A-B)**. CD4 frequencies and numbers trended downward but did not reach statistical significance between treatment groups **(Fig S4C)**. FTY720 also reduced B cell counts in the blood of E7+FTY720 treated groups compared to the E7 control **(Fig S4D)**. γδ T cell counts did not change with treatment **(Fig S4E)**.

**Figure 5:**
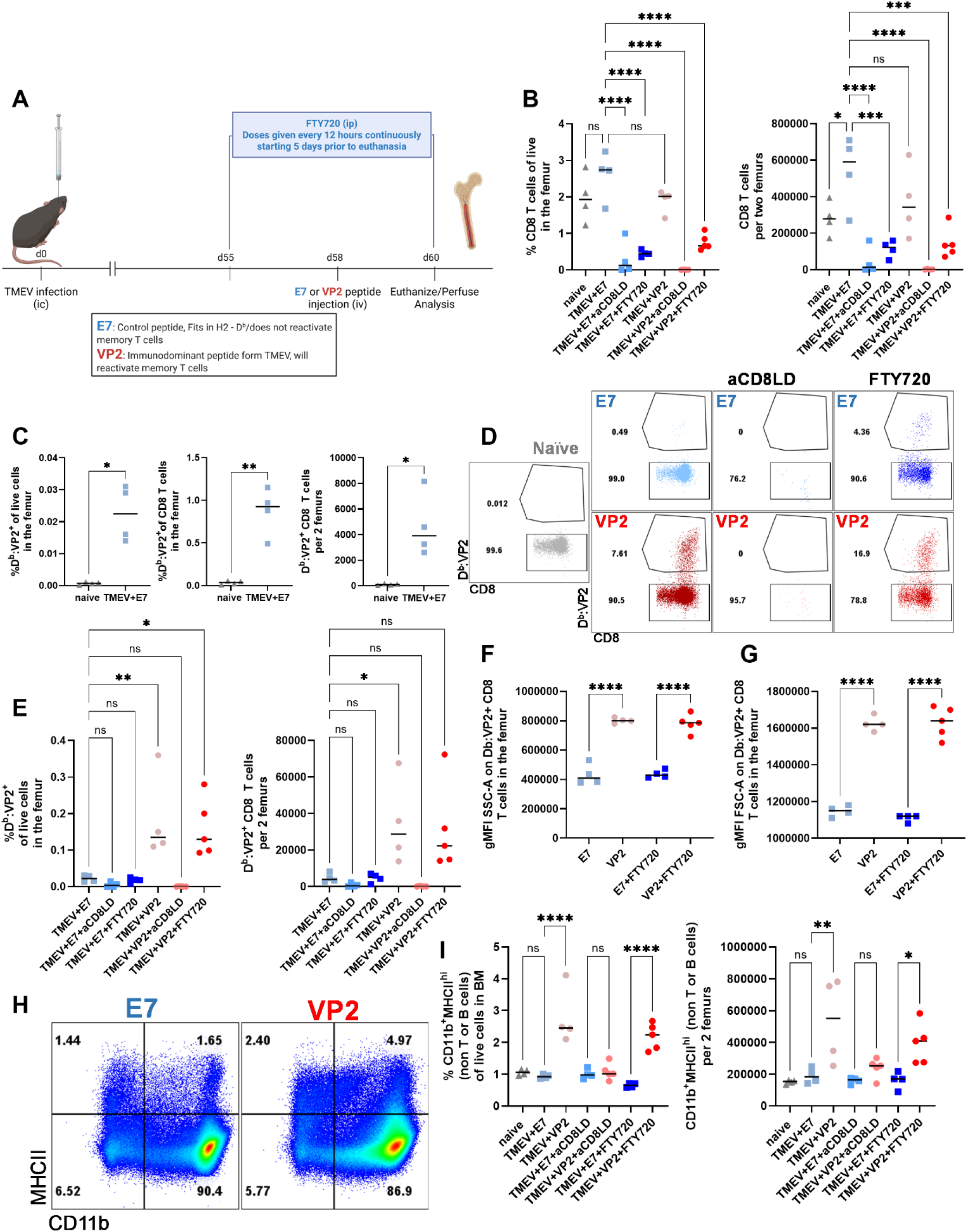
Antigen-specific CD8 T cells generated within the bone marrow following neurotropic TMEV infection become durable memory and can be reactivated upon antigen reencounter. **A)** Experimental design is shown. Mice were intracranially infected with TMEV and allowed to rest for 55 days before FTY720 and/or anti CD8 treatment (low dose) according to the schematic. E7 or VP2 peptides were given iv two days before euthanasia. Femoral bone marrow was analyzed. **B)** Frequencies and counts of CD8 T cells in the bone marrow were reduced in FTY720 and aCD8LD groups, as expected. **C)** Compared to the naïve (never infected) mice, previously infected quiescent mice have significantly higher frequencies and counts of D^b^: VP2+ CD8 T cells indicating formation of durable antigen-specific memory within the femur bone marrow niche 60 days after brain viral infections. **D-E)** Antigen-specific CD8 T cells within the bone marrow are found in previously infected mice and expand upon antigen reencounter in VP2 and VP2+FTY720 groups. In aCD8LD treated groups, all CD8s are significantly depleted. **F-G)** gMFI of SSC-A (Side Scatter Area) and FSC-A (Forward Scatter Area) on D^b^: VP2+ CD8 T cells significantly increases in VP2 and VP2+FTY720 groups compared to the respective E7 controls indicating blasting. **H-I)** Reactivation of the antigen-specific CD8 T cells is associated with an increase in frequencies and counts of CD11b+ MHCII+ cells within the bone marrow in VP2 and VP2+FTY720 groups compared to the respective E7 controls. This is while the CD8 depleted groups had no change in their CD11b+ MHCII+ populations indicating the myeloid influx is CD8 T cell dependent. In **B** and **E**, a one-way ANOVA followed by Dunnett’s multiple comparisons test was performed and all groups were compared to the E7 control. In **C**, Welch’s t-test was performed. **D** and **E** show representative flow plots. In **D**, cells within the gated CD8 T cells population are shown. In **H**, CD11b and MCHII expression on TCRβ-B220-(non-T and non-B cells) cells are shown. In **F**, **G**, and **I**, a one-way ANOVA followed by Šídák’s multiple comparisons test was performed comparing the selected pairs. Data are shown as individual mice with mean and SD. Comparisons made are shown on the graph. ns = P > 0.05, * = P ≤ 0.05, ** = P ≤ 0.01, *** = P ≤ 0.001, and **** = P ≤ 0.0001.

Having verified that FTY720 and CD8 depleting antibodies both significantly reduced CD8 T cell counts in the blood, we next evaluated the antigen-specific CD8 T cell responses in the bone marrow. First, we quantified CD8 T cell frequencies and counts within the femoral bone marrow. Compared to the E7 control groups, both FTY720 and low dose anti CD8 depletion reduced CD8 T cell counts and frequencies in treated E7 or VP2 groups **(Fig 5B)**. Total cells recovered from all bone marrow samples were similar **(Fig S4F)**. Minor differences in total CD4 frequencies and counts were also observed when comparing previously infected E7 groups to E7+aCD8LD, E7+ FTY720, VP2+aCD8LD, or VP2+FTY720 groups **(Fig S5G)**.

We next evaluated if antigen-specific CD8 T cells within the bone marrow are durably stable 60 days after TMEV infection. To answer this question, we compared frequencies and counts of D^b^: VP2+ CD8 T cells in the femurs of naïve (never infected) mice to those previously infected but not reactivated (TMEV+E7) group and found significant numbers of virus antigen-specific CD8 T cells in the femur 60 days post-infection **(Fig 5C**, from left to right, % D^b^: VP2+ CD8 T cells of live, D^b^: VP2+ CD8 T cells of total CD8 T cells, and absolute counts of D^b^: VP2+ CD8 T cells). Using representative flow plots of gated CD8 T cells, we also verified that compared to the naïve group, TMEV+E7, and TMEV+ FTY720 groups both contained detectible frequencies of antigen-specific CD8 T cells in the femoral bone marrow **(Fig 5D-E)**. As expected, CD8 depleting antibodies depleted the majority of the CD8 T cells including the antigen-specific CD8 T cells from the bone marrow of TMEV+E7+aCD8LD and TMEV+VP2+aCD8LD treated groups **(Fig 5D-E)**. We next evaluated the extent to which VP2 administration reactivates bone marrow resident antigen-specific CD8 T cells. VP2 administration reactivated antigen-specific CD8 T cells within the bone marrow and their responses were marked by a rapid increase in frequencies and numbers of D^b^: VP2+ CD8 T cells in TMEV+VP2 groups **(Fig 5D-E)**. FTY720 did not prevent the reactivation of D^b^: VP2+ CD8 T cells in the bone marrow as their frequencies still increased in TMEV+VP2+FTY720 groups compared to the E7 control groups **(Fig 5D-E)**. Counts of D^b^: VP2+ CD8 T cells in the bone marrow of TMEV+VP2+FTY720 groups trended to increase as well compared to controls, though it did not reach statistically significant results, likely due to the reduction in total CD8 T cell counts **(Fig 5E)**. We next evaluated if reactivation of antigen-specific memory CD8 T cells in the bone marrow 60 days post intracranial infection was associated with the blasting phenotype in antigen-specific CD8 T cells. To evaluate this, we compared the geometric MFI of side scatter and forward scatter (area) parameters between previously infected E7 and VP2 and E7+FTY720 and VP2+FTY720 groups. We determined that D^b^: VP2+ CD8 T cells in the reactivated mice (VP2 groups) had significantly higher side and forward scatters indicating T cell activation and blasting **(Fig 5F-G)**.

Finally, we evaluated the extent antigen-specific CD8 T cell reactivation in the bone marrow was associated with an influx of inflammatory myeloid cells into the bone marrow niche, a feature we also observed in the brain following reactivation^17^. We compared previously infected quiescent bone marrow to marrow of mice administered VP2 peptide 2 days prior to restimulate antigen-specific CD8 memory T cells. In mice receiving VP2 peptide, we found significantly higher frequencies and numbers of CD11b+ MHCII+ myeloid cells **(Fig 5H-I)**. The influx of these likely inflammatory myeloid cells within the bone marrow was observed in VP2 and VP2+FTY720 treated groups but was abrogated in the CD8 depleted groups **(Fig 5I)**. These data indicate that CD8 T cells directly lead to an influx of MHCII+ myeloid cells within the bone marrow following antigen-specific reactivation 60 days post-infection, a time point consistent with memory formation **(Fig 5I)**. In summary, we determined that 60 days post brain viral infections, durable antigen-specific CD8 T cells are found within the bone marrow. These memory CD8 T cells can be reactivated upon reencounter with their cognate antigen, expand in response to reactivation, and undergo blasting. Reactivation of antigen-specific memory CD8 T cells in the bone marrow is not dependent on the influx of new CD8 T cells from the blood or SLO as FTY720 did not block CD8 T cell reactivation, expansion, or blasting. Finally, antigen-specific reactivation of CD8 T cells in the bone marrow is associated with increased influx of MHCII+ myeloid cells. This influx is directly caused by CD8 T cells as their depletion abrogates the myeloid response.

### Virus antigen-specific CD8 T cells generated in the bone marrow following neurotropic TMEV infection are durable 200 days post-infection and can still be reactivated upon antigen reencounter

Mice were intracranially infected with TMEV and 15-28 weeks were allowed to pass. Then 5 days before euthanasia, FTY720 or CD8 depletion was initiated as before. In addition to the low dose CD8 depletion group (aCD8LD), we also added high dose CD8 depletion groups (aCD8HD) in these experiments to better deplete CD8 T cells in all tissues. E7 or VP2 peptides were intravenously delivered 2 days before euthanasia **(Fig 6A)**. When comparing E7 groups, CD8 depletion strategies reduced frequencies and counts of CD8 T cells within the femoral bone marrow **(Fig 6B)**. When comparing VP2 groups, FTY720, aCD8LD, and aCD8HD treatment reduced CD8 T cell frequencies and counts as expected and previously seen **(Fig 6B)**. Frequencies and counts of CD4 T cells in the bone marrow remained largely comparable in the femoral bone marrow **(Fig S5 A)**. Similar to day 60, previously infected quiescent (E7) mice had significantly higher numbers of D^b^: VP2+ CD8 T cells in the femoral bone marrow compared to naïve never infected controls, indicating durable memory within the bone marrow even 200 days after a brain infection **(Fig 6C)**. Frequencies of D^b^: VP2+ CD8 T cells in the femoral bone marrow increased between E7 and VP2 groups indicating reactivation **(Fig 6D**, and **Fig S5B)**. Counts of D^b^: VP2+ CD8 T cells in the femoral bone marrow did not reach statistically different results between E7 and VP2 groups **(Fig 6D)**. As expected, CD8 T cell depletion, significantly reduced frequencies and numbers of D^b^: VP2+ CD8 T cells in the femoral bone marrow **(Fig 6D**, and **Fig S5B)**. We next evaluated if D^b^: VP2+ CD8 T cells in the femoral bone marrow blast following reactivation and antigen reencounter via measuring gMFI of SSC-A and FSC-A. We found that in two independent experiments, gMFI of SSC-A and FSC-A on D^b^: VP2+ CD8 T cells in the femoral bone marrow significantly increased in VP2 and VP2+FTY720 treated compared to E7 and E7+FTY720 groups **(Fig 6E-F**, and **Fig S5C-D)**. It is worth mentioning that 3-minutes prior to euthanasia, mice received an intravenous injection of fluorescently labeled αCD45 antibody to label immune cells in or near vasculature. The overwhelming majority of D^b^: VP2+ CD8 T cells in the femoral bone marrow were iv- indicating tissue residency in E7/E7+FTY and VP2/VP2+FTY720 groups **(Fig S5E)**.

**Figure 6:**
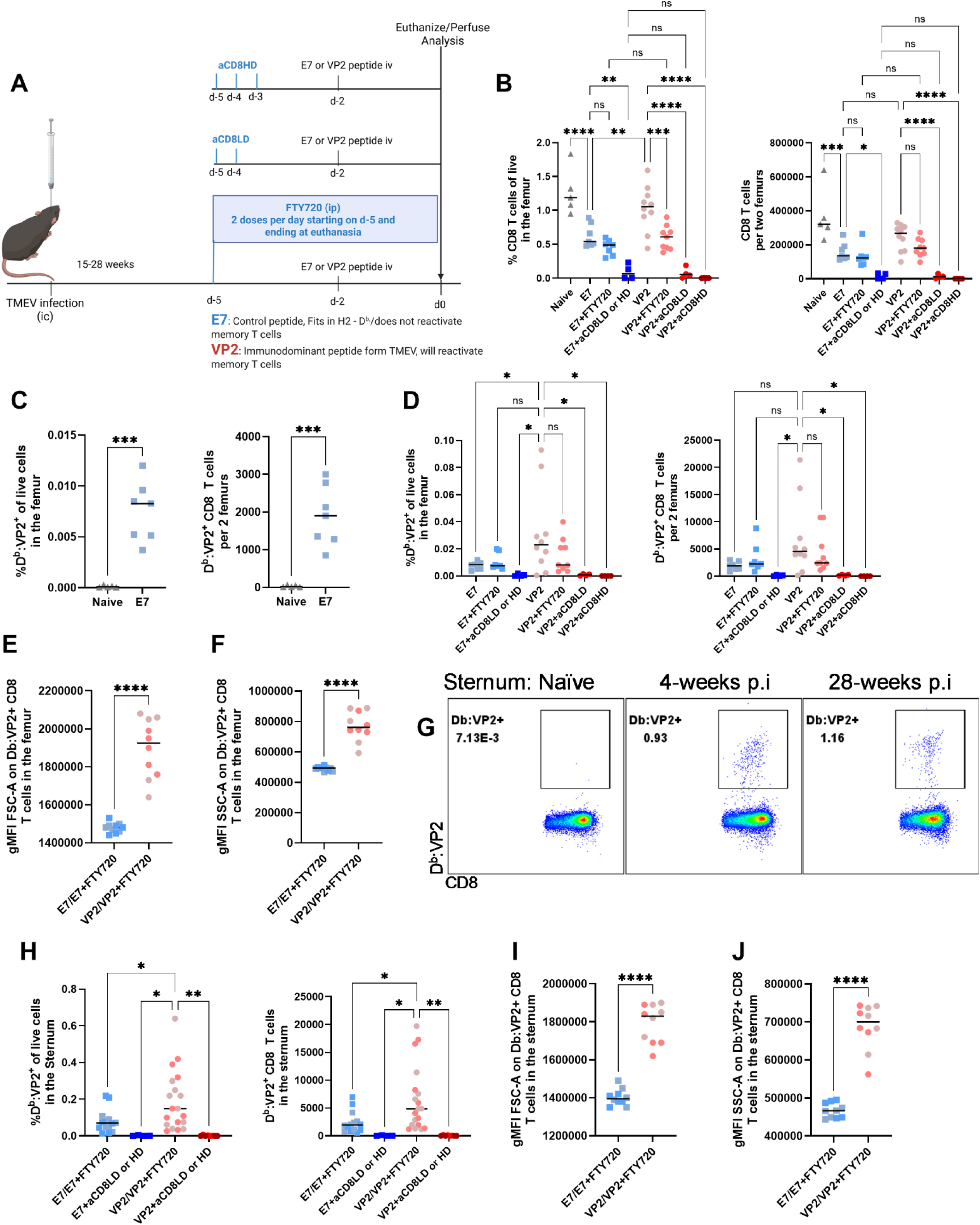
Virus antigen-specific CD8 T cells generated following neurotropic TMEV infection remain within the bone marrow niche long-term and can be reactivated upon systemic antigen reencounter. **A)** Experimental design and treatment schedule are shown. Mice were intracranially infected with TMEV and allowed to rest 15-28 weeks. One set of experiments was performed at 15 weeks and one set at 28 weeks post infection as such experiments were performed twice. Similar results were obtained and hence results shown are pooled from these two independent experiments. FTY720 and aCD8LD and aCD8HD treatment schedules are as shown and described in the results. E7 or VP2 peptides were given intravenously two days before euthanasia. 3-minutes before euthanasia, mice received an intravenous injection of fluorescently labeled αCD45 antibody. **B)** Frequencies and counts of total CD8 T cells in the femoral bone marrow are shown and demonstrates efficacious depletion (as expected) in the depleting conditions. Since we found no differences between control E7 aCD8LD and aCD8HD groups, these groups are combined on the graph. **C)** Frequencies and numbers of antigen-specific CD8 T cells are compared between naïve and previously infected mice. Antigen-specific CD8 T cells in the femoral bone marrow are durable post brain viral infections. **D)** Frequencies and numbers of antigen-specific CD8 T cells in the femoral bone marrow are compared between groups. **E-F)** Reactivation of antigen-specific CD8 T cells within the bone marrow results in blasting of antigen-specific T cells in the femurs measurable by increased gMFI of FSC-A and SSC-A on these cells. **G)** In the sternal marrow, antigen-specific CD8 T cells against TMEV are found at early and late times post brain infection. Representative flow plot is shown (gated CD8 T cells). **H)** Frequencies of antigen-specific CD8 T cells in the sternal bone marrow increase between E7 and VP2 groups and are depleted, as expected, in the depleted groups. There were no differences between E7 and E7+FTY720 and VP2 and VP2+FTY720 groups and as such these groups were combined on the graph. In **E-F**, and **H-J**, the symbol color represents treatment groups and follows previous graphs with FTY720 treated mice being shown by darker colors. **I-J)** Antigen-specific CD8 T cells within the sternal bone marrow blast upon antigen reencounter which is measurable by increased gMFI of FSC-A and SSC-A on these cells. In **B**, **D**, and **H**, a one-way ANOVA with Šídák’s multiple comparisons test or Dunnett’s multiple comparisons test was performed comparing between selected groups. Comparisons are shown on the graph. In **C**, **E-F**, and **I-J**, a Welch’s t test was performed comparing the two groups. Color scheme in pooled groups (**E-F, H-J**) follows individual groups (**B**, and **D**). Data are shown as individual mice with mean and SD. Comparisons made are shown on the graph. ns = P > 0.05, * = P ≤ 0.05, ** = P ≤ 0.01, *** = P ≤ 0.001, and **** = P ≤ 0.0001.

### Sternal bone marrow maintains populations of virus antigen-specific memory CD8 T cells that can be reactivated upon antigen reencounter long after TMEV infection is cleared

We next evaluated antigen-specific CD8 T cell responses in the sternal bone marrow. Similar to previous results, antigen-specific CD8 T cells were found in previously infected sternal bone marrows at early (4 weeks post infection) and late memory (28 weeks post infection) time points **(Fig 6G)**. As expected, CD8 T cell depletion reduced frequencies and counts of CD8 T cells and D^b^: VP2+ CD8 T cells in the sternal bone marrow **(Fig S5F**, and **Fig 6H**). Frequencies and counts of CD4 T cells within the sternal bone marrow did not change between groups **(Fig S5G)**. Compared to naïve uninfected mice, previously infected mock E7 peptide treated controls had a durable population of D^b^: VP2+ CD8 T cells in the sternal bone marrow **(Fig S5H)**. D^b^: VP2+ CD8 T cells in the sternal bone marrow were reactivated upon antigen exposure via expansion and blasting **(Fig 6 H-J, and Fig S5I-J)**. Finally, as seen previously and reported in femurs **(Fig S5E)**, D^b^: VP2+ CD8 T cells in the sternal bone marrow were iv- indicating that the majority of antigen-specific CD8 T cells are tissue resident **(Fig S5K)**. D^b^: VP2+ CD8 T cells in the femur and sternal bone marrows therefore form durable memory up to 28 weeks post brain infection and can reactivate independently of the blood and SLO T cells upon cognate antigen reencounter.

### Antigen-specific reactivation of CD8 T cells within the femoral bone marrow is associated with the dysregulation of hematopoietic stem and progenitor cells

Thus far, we demonstrated that brain viral infections generate an acute and memory antigen-specific T cell response within the bone marrow niche that is durable and can be reactivated (at the memory time-point) upon antigen reencounter. We next sought to determine the impact of antigen-specific memory CD8 T cell reactivation on the stem cell niche. In the experiments described in **Fig 6A**, we evaluated lineage-Sca1+ ckit+ (LSK) cell populations. We found that in femurs of naïve and previously infected but quiescent mice, low levels of LSK cells are found **(Fig 7A** comparing naïve to E7 groups). However, antigen-specific reactivation of D^b^: VP2+ CD8 T cells resulted in a massive dysregulation of LSK cells in the femoral bone marrow, exhibited by expansion in the absence of active viral infection or danger signals **(Fig 7A** comparing E7 to VP2 and VP2+FTY720 groups). FTY720 treatment did not block the LSK dysregulation observed following reactivation of memory CD8 T cells through VP2 peptide administration **(Fig 7A)**. When we evaluated quiescent LSKs by quantifying CD38 low LSK populations, we found similar results. Antigen-specific reactivation results in the expansion of CD38low LSK cells in VP2 or VP2+FTY720 treated groups compared to E7 controls **(Fig 7B**, and **Fig S6A)**. This expansion of LSK cells as a result of antigen-specific CD8 T cell reactivation is not explained by changes in the femoral bone marrow cellularity as comparable recovered total cells were seen in all groups **(Fig S6B)**. We next evaluated if the LSK dysregulation was CD8 T cell dependent or a result of VP2 peptide administration through comparing CD8 T cell depleted groups (VP2+aCD8LD and VP2+aCD8HD).

**Figure 7:**
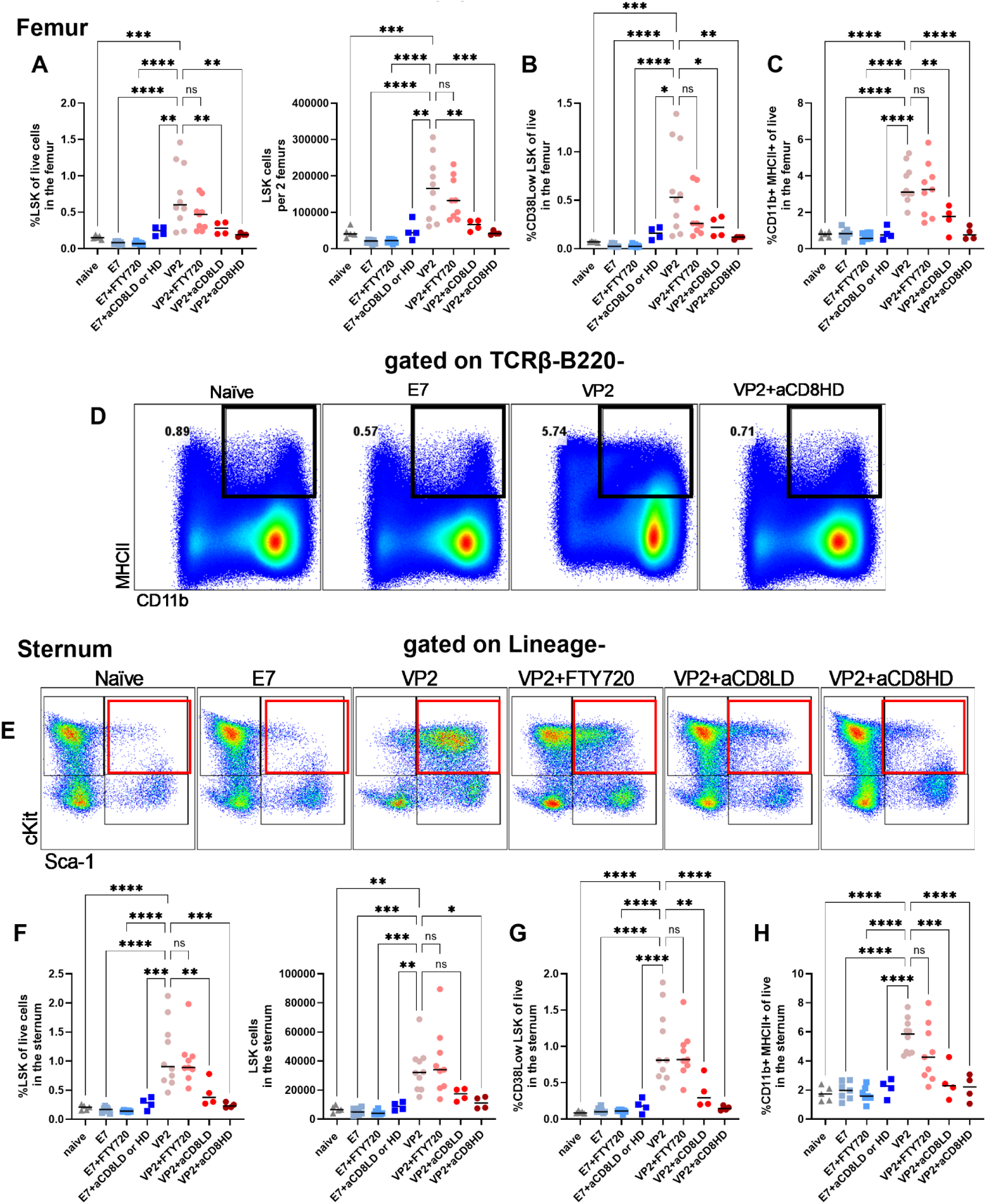
Antigen-specific reactivation of memory CD8 T cells within the bone marrow impacts stem cell populations. Experimental design is as previously described in **Fig 6A**. The stem cell populations from the same mice analyzed in Fig 6 are shown here. **A)** Frequencies and number of LSK (Lineage-, Sca-1+, cKit+) cells increase in VP2 and VP2+FTY720 treated mice compared to controls. CD8 T cell depletion abrogates this LSK dysregulation. Lineage negative was defined as single cells, live, TCRβ-, TCRγδ-, B220-, Ly6C-, Ly6G-, NK1.1-, CD11c-, CD11b-, F480-, CD45iv-. **B)** Frequencies of CD38 LSK cells are shown and follow the same dysregulation pattern seen in VP2 and VP2+FTY720 treated groups. CD8 depletion abrogates this LSK dysregulation following antigen-specific memory CD8 T cells dysregulation. **C-D)** Concurrent with antigen-specific CD8 T cells reactivation and LSK dysregulation, we also observed an influx of CD11b+ MHCII+ inflammatory myeloid cells into the femoral bone marrow. In **D**, black boxes designate the population quantified. The representative flow plots are showing the TCRβ-B220-populations. **E-F)** Similar to the femur, sternal bone marrow also responds to the antigen-specific memory CD8 T cell reactivation by LSK expansion. LSK expansion following memory CD8 T cell reactivation occurs in VP2 and VP2+FTY720 treated groups compared to controls but is abrogated in the CD8 T cell depleted groups. In **E**, red boxes show the LSK population. Representative flow plots are gated on lineage negative populations. **G)** CD38 low LSK cells are also dysregulated following antigen-specific CD8 T cell reactivation in the sternal bone marrow in a CD8 dependent manner. **H)** In the sternum, antigen-specific memory CD8 T cell reactivation induces an influx of CD11b+ MHCII+ cells. This influx is abrogated when CD8 T cells are depleted. In **A-C** and **F-H**, a one-way ANOVA with Dunnett’s multiple comparisons test was performed comparing between the selected groups. Comparisons made are shown on the graph. Data are shown as individual mice with mean and SD. ns = P > 0.05, * = P ≤ 0.05, ** = P ≤ 0.01, *** = P ≤ 0.001, and **** = P ≤ 0.0001.

Interestingly, CD8 T cell depletion abrogated the LSK and CD38 low LSK dysregulation in VP2-injected mice indicating that antigen-specific reactivation of memory CD8 T cells directly induces LSK dysregulation within the femoral bone marrow **(Fig 7A-B**, and **Fig S6A)**.

Antigen-specific reactivation of CD8 T cells also led to an increase in CD11b+ MHCII+ cells within the femoral bone marrow **(Fig 5H)**. We therefore evaluated if this population also infiltrated upon antigen reencounter at the late memory timepoint (15-28 weeks post infection). We observed an increase in CD11b+MHCII+ cells in VP2 and VP2+FTY720 treated groups compared to naïve or E7 controls **(Fig 7C-D**, and **Fig S6C)**. The influx in CD11b+ MHCII+ cells within the femoral bone marrow following antigen-specific reactivation was abrogated when CD8 T cells were depleted **(Fig 7C**, and **Fig S6C)**. Hence in the femur, antigen-specific CD8 T cell reactivation resulted in an increase in LSK populations concurrent with an influx of inflammatory myeloid cells. LSK dysregulation and myeloid cell influx into the femoral bone marrow was CD8-dependent.

### Virus antigen-specific reactivation of memory CD8 T cells within the sternal bone marrow is associated with the dysregulation of hematopoietic stem and progenitor cells

Finally, we evaluated LSK counts, CD38low LSK numbers, and the influx of inflammatory myeloid cells into the sternal bone marrow niche following antigen-specific CD8 T cell reactivation at the memory time points after brain viral infections. Similar to the femoral bone marrow results, antigen-specific CD8 T cell reactivation resulted in a significant increase in LSK cells, and CD38low LSK cells. FTY720 treatment did not change LSK dysregulation following antigen-specific CD8 T cell reactivation in the sternum indicating blood and SLO T cells do not contribute to this phenomenon **(Fig 7E-G, and Fig S6D)**. Importantly, LSK dysregulation was exclusively dependent on CD8 T cells alone as their depletion completely abrogated the LSK and CD38low LSK dysregulation following antigen-specific memory CD8 T cell reactivation in the bone marrow **(Fig 7E-G, and Fig S6D)**. Similar to the femur, the total numbers of T cells recovered from the sternal bone marrow across conditions were similar indicating that the LSK dysregulation is not simply due to a change in the sternal bone marrow cellularity **(Fig S6E)**. The influx in MHCII+ myeloid cells within the bone marrow was also observed in the sternum of VP2 and VP2+FTY720 treated mice compared to controls, while CD8 depletion abrogated this increase **(Fig 7H, and Fig S6F)**.

In summary, antigen-specific CD8 T cells following brain viral infections remain in the bone marrow and can be reactivated long after the infection is cleared. This reactivation occurs independently of the circulating CD8 T cells as it can occur despite FTY720 treatment. Importantly, antigen-specific memory CD8 T cell reactivation greatly impacts LSK expansion and influx of inflammatory myeloid cells into the bone marrow. These hematopoietic effects are CD8 T cell dependent as CD8 depletion prior to peptide antigen-specific reactivation completely abrogates the LSK dysregulation and CD11b+ MHCII+ influx into the bone marrow. We did not observe differences between the femoral or sternal marrow responses in terms of antigen-specific CD8 reactivation and LSK dysregulation or myeloid influx indicating systemic antigen-specific reactivation can equally impact multiple sources of the bone marrow.

## Discussion

In this study, we sought to understand whether brain viral infections lead to the generation and maintenance of antigen-specific CD8 T cells within the bone marrow compartment plus determine the consequences of antigen-specific CD8 T cell reactivation within the bone marrow niche. Our studies comprehensively evaluated priming, memory generation, and reactivation of antigen-specific CD8 T cells within the bone marrow following neurotropic viral infection of the brain. We also sought to evaluate the impact of acute brain insults, including infection with the neurotropic virus TMEV on the bone marrow niche. We found that acute brain infection with TMEV generates a population of virus antigen-specific CD8 T cells within the bone marrow compartment **(Fig 1)**. These antigen-specific CD8 T cells do not depend on major T cell trafficking through the blood and SLO as FTY720 treatment (which effectively blocks T cell exit out of the SLO into the blood circulation) did not impact frequencies of antigen-specific CD8 T cells in the bone marrow following TMEV infection **(Fig 2)**. Importantly, the generation of antigen-specific CD8 T cells within the bone marrow compartment was not a unique feature of TMEV infection, or D^b^: VP2 immune responses, as delivery of Ovalbumin admixed with Poly I:C into the brain similarly generated an antigen-specific CD8 T cell (K^b^: SIINFEKL) response within the bone marrow **(Fig 4)**. Importantly, we demonstrated that antigen-specific CD8 T cell responses in the bone marrow following brain infection generated and established durable memory responses that lasted long-term and could be reactivated upon antigen reencounter **(Fig 4-6)**. Reactivation of antigen-specific CD8 T cells in the bone marrow led to a CD8-dependent influx in inflammatory myeloid cells and dysregulation of HSPCs within the bone marrow **(Fig 7)**.

### T cells have been found in the bone marrow under normal conditions and following systemic insults

Our studies are in line and add to the body of literature regarding the functions of T cells in the bone marrow under normal conditions^58^, as well as sequestration of T cells into the bone marrow following brain cancer^26^ or systemic infections^7^. Under normal conditions, in mice and humans, populations of naïve and memory CD4 and CD8 T cells have been found in the bone marrow compartment. In humans, antigen-specific T cells against common viral infections including CMV and childhood vaccinations (tetanus, measles, and rubella) have been evaluated and quantified^58–73^. In fact, a specific population of memory CD8 T cells seems to prefer the bone marrow for homeostatic proliferation^74^. These studies were seminal in describing T cells in the healthy bone marrow and indicating that both in mice and humans, peripheral infections and/or vaccinations can induce antigen-specific T cell responses within the bone marrow niche.

In mice, it has been shown that antigen-specific CD8 and CD4 T cells are found in the bone marrow following systemic viral and bacterial infections, including LCMV, influenza, VSV, and *Listeria monocytogenes* infection ^72, 73, 75, 76^. Additionally in mice, antigen-specific CD8 T cells against LCMV were tracked in the bone marrow and demonstrated to be not only be present but also functionally retain their cytotoxic capacity^72^. The homing and establishment of these cells into the bone marrow, while still an emerging field, is likely dependent on CXCR4, CCL12, IL-7, IL-15, and CD69 ^58, 77–79^. Interestingly, in one recent study, authors found that influenza infections impacted HSPCs in the bone marrow and skewed megakaryocyte generation possibly contributing to abnormal clotting seen post influenza infection^80^. While this study did not directly evaluate the contribution of antigen-specific CD8 T cells in the bone marrow, their results are in line with ours and could suggest involvement of CD8 T cells in HSPCs dysregulation resulting in the eventual dysregulation of megakaryocyte production.

### Antigen-specific CD8 T cell responses following infections can be primed in the bone marrow by APCs bringing antigens from the brain into the bone marrow niche

In our experiments in which AF488-labeled OVA was directly delivered into the brain, we demonstrated that antigen delivery into the brain was concurrent with antigen pick up by APCs, including myeloid cells expressing CD11b+ MHCII+ and CD103 **(Fig 3)**. AF488 and the SIINFEKL peptide bound to H2-K^b^ were both detectible in the brain infiltrating myeloid APCs. Interestingly, these populations of APCs were also present in the bone marrow and had detectible antigen (AF488) within them, indicating their origin before trafficking was the brain. We contend these myeloid APCs with migratory potential have the capacity to *in situ* prime CD8 T cells within the bone marrow following acute brain insults. Interestingly, CD8 T cells can migrate to the bone marrow where they can be found in tight proximity to CD11c+ dendritic cells, monocytes, macrophages, and neutrophils; these antigen presenting cells in the bone marrow are generally thought to process and present antigens to prime naïve CD8s ^81–84^. As such, our study adds depth to the above body of previous research while also contributing the additional “brain-specific” context which was previously understudied.

### T cells in the bone marrow are relevant to CNS cancers

In the context of brain cancer, it was shown that in mice with an aggressive form of brain cancer, glioblastoma, T cells become sequestered within the bone marrow compartment^26, 27^. This T cell sequestration in the context of brain cancer is thought to be a feature of systemic immunosuppression^85^. However, in these brain cancer studies, total T cells, and not antigen-specific T cells were evaluated. Hence, whether tumor antigen-specific T cells accumulate within the marrow is yet to be determined and is an active area of investigation in our laboratory. In subcutaneous tumor models, however, a recent study evaluated the function of bone marrow CD8 T cells against tumor infiltrating lymphocytes (TILs) in the B16-OVA implanted mice and found bone marrow resident antigen-specific CD8 T cells to be more stem-like than TILs^86^. This study showed stable numbers of OVA-specific CD8 T cells in the bone marrow and upon adoptive transfer, they performed better than TILs in terms of persistence, tumor infiltration, and lack of exhaustion^86^. Although less relevant to our studies, in patients with a severe form of aplastic anemia, T cells with a TRM-like phenotype with high capacity to produce cytokines including IFNγ were also found in the bone marrow and are thought to assist in HSPC destruction^87^. Though this study was done in the context of cancers of the bone marrow, TRM cytokine productions could mechanistically explain our results presented in **Fig 7**, in which CD8 TRM reactivation led to an increase in myeloid cells and LSK cells. Mechanistic insight into bone marrow resident T cell cytokine production and responses are also amongst active areas of investigation within our lab.

### Concluding remarks and impact

We demonstrated that the bone marrow niche responds to brain insults by generating populations of antigen-specific CD8 T cells that can generate durable memory within the bone marrow niche. These TRMs can be reactivated upon reencounter and expand and initiate blasting processes. TRM reactivation also leads to a significant influx of inflammatory myeloid cells in the bone marrow (even in the absence of an actual viral infection or danger signals). Concurrently, TRM reactivation in the bone marrow leads to increased LSK numbers, indicating that T cells in the bone marrow directly interact with and/or impact HSPCs. Previous studies discussed above determined that peripheral infections and vaccinations, in both mice and humans, also lead to the generation and maintenance of populations of antigen-specific T cells in the bone marrow. Given our results are in line with the literature, we contend antigen-specific T cells in the bone marrow are generated in response to local and systemic insults and can impact the complicated process of hematopoiesis. Our ongoing studies will investigate mechanisms through which T cells impact HSPCs. Unresolved questions include understanding how repeated reactivation of bone marrow resident antigen-specific T cells impacts hematopoiesis and shapes the future HSPC and/or plasma cell responses within the niche. Similarly, whether these functional stem-like bone marrow resident antigen-specific CD8 T cells can be harnessed to eradicate cancers in various tissues, including the brain is yet to be determined. Theoretically, releasing these sequestered populations of antigen-specific T cells from the bone marrow could improve anti-tumor immunity, as these immune cells have favorable proliferative, maintenance, and cytokine release upon engaging antigens. These studies therefore provide additional understanding of bone marrow T cell responses following brain insults and provide irrefutable evidence that bone marrow TRM responses impact other cells within the bone marrow niche.

## Acknowledgments

We would like to thank Fang Jin, Michael J Hansen, Mark Maynes, and Frances Rangel for their help with our experiments and scientific discussions. We thank the NIH tetramer facility for providing D^b^: VP2 tetramer. We acknowledge Biorender.com for help with figure generation.

## Author contributions

DMAW, KA, RAR, CKP, EMM, SEW, CEF, AMHM, VJ, KEB, PKN performed the experiments and provided intellectual contributions. KA, KMH, AJJ, DMAW, and KEB conceptualized the project, provided intellectual contributions and mentored other authors. KA, DMAW, RAR, KMH, AJJ, and VJ wrote the manuscript. DMAW, RAR, and KA analyzed the data. KA made final figures and wrote the first draft of the manuscript. All authors read and edited a version of the manuscript. MK and GAG provided financial and salary support for KMH and KEB (respectively) and intellectually contributed to the manuscript. VC intellectually contributed, read and helped edit the manuscript. AJJ and KA were financially responsible for all mice, reagents, and staff salaries.

## List of abbreviations

APC: Antigen presenting cells
VP2: Viral antigen 2
VP2 peptide: VP2_121-130_ peptide
TMEV: Theiler’s murine encephalomyelitis virus
LCMV: Lymphocytic choriomeningitis virus
VSV: Vesicular stomatitis virus
S1P: Sphingosine-1-phosphate
BM: Bone marrow
TRM: Tissue resident memory T cells
HSPCs: hematopoietic stem and progenitor cells
DC: Dendritic cells
SD: Standard deviation
OVA: Ovalbumin
Poly I:C: Polyinosinic:polycytidylic acid
LSK: Lineage-Sca1+ cKit+
RT: Room temperature
SLO: secondary lymphoid organs
Min: Minutes (when mentioned in the methods section and is referring to organ processing and staining), Minimum when showing heatmaps
Max: Maximum
°C: Degrees Celsius
Fig: Figure
iv: Intravenous
RCF: relative centrifugal force
g: grams
× g: Gravitational force
RPMI: Roswell Park Memorial Institute media
PBS: Phosphate Buffered Saline

---

**Figure S1:**
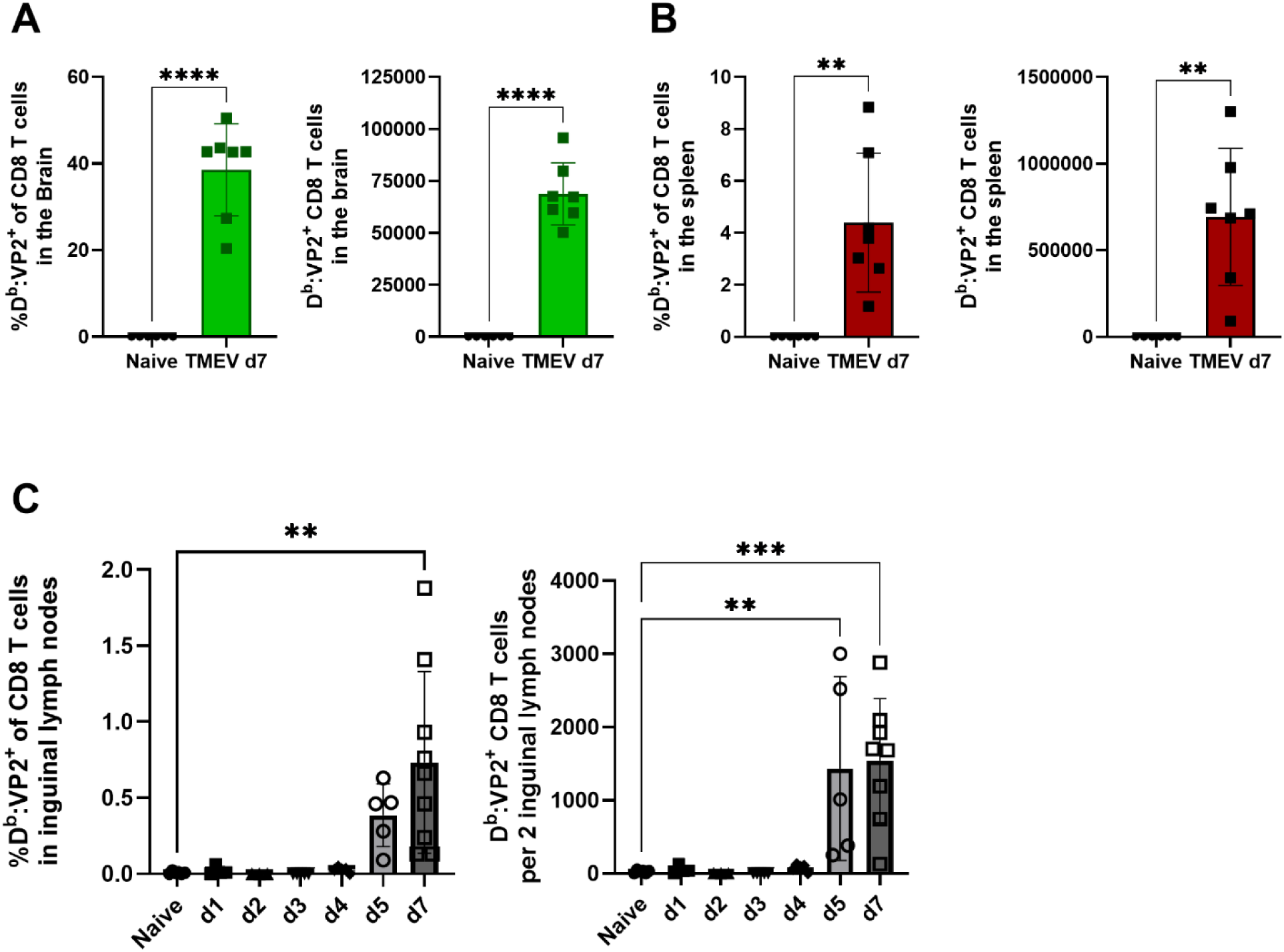
Virus antigen-specific CD8 T cells accumulate in the secondary lymphoid organs and brain following TMEV infection. Experimental design is as shown in Figure 1A. **A-C)** Frequencies and absolute counts of D^b^: VP2+ CD8 T cells are shown in the brain **(A)** and the spleen **(B)** and the inguinal lymph nodes **(C)** of either uninfected or TMEV infected mice at various times post-infection. In **A-B**, an unpaired two-tailed Welch’s t-test was performed. In **C**, a Kruskal-Wallis mixed effects model (REML) with the post-hoc Dunn’s multiple comparisons test was performed comparing naïve to each time point. Data are shown as individual mice. Mean and SD are shown. ns = P > 0.05, * = P ≤ 0.05, ** = P ≤ 0.01, *** = P ≤ 0.001, and **** = P ≤ 0.0001.

**Figure S2:**
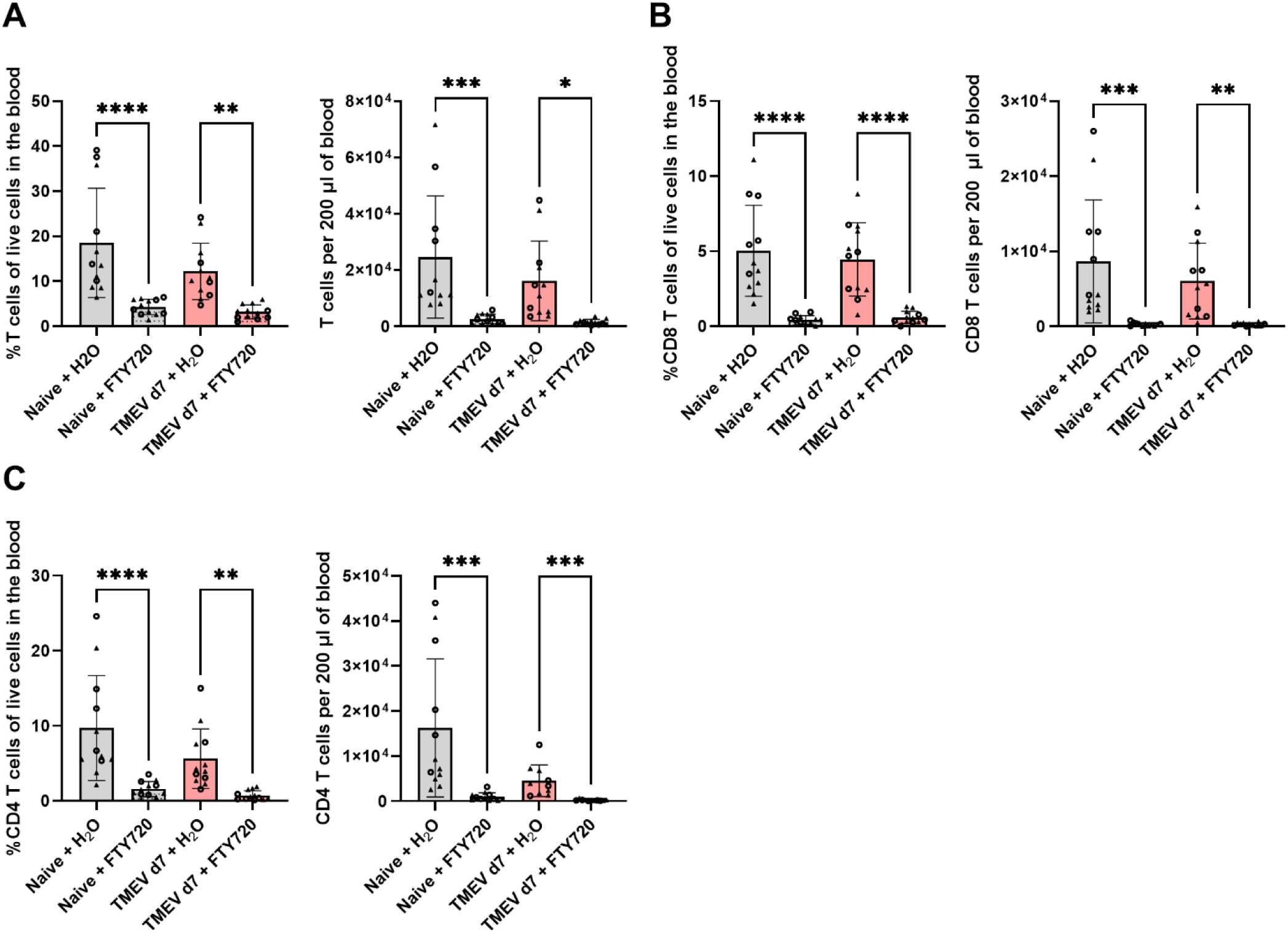
FTY720 significantly reduces circulating T cells in the blood. **A-C)** Frequencies and numbers of total T cells, CD8 T cells, and CD4 T cells are quantified in the blood. The Shapiro Wilk’s normality test was performed. For normally distributed data, a one-way ANOVA was used and for non-normally distributed data a Kruskal-Wallis test was performed. Holm-Šídák’s post-hoc analysis was performed following the ANOVA while Dunn’s multiple comparisons test was performed post Kruskal-Wallis analysis. Data are shown as individual mice. Mean and SD are shown. Comparisons made are shown on the graph. ns = P > 0.05, * = P ≤ 0.05, ** = P ≤ 0.01, *** = P ≤ 0.001, and **** = P ≤ 0.0001.

**Figure S3:**
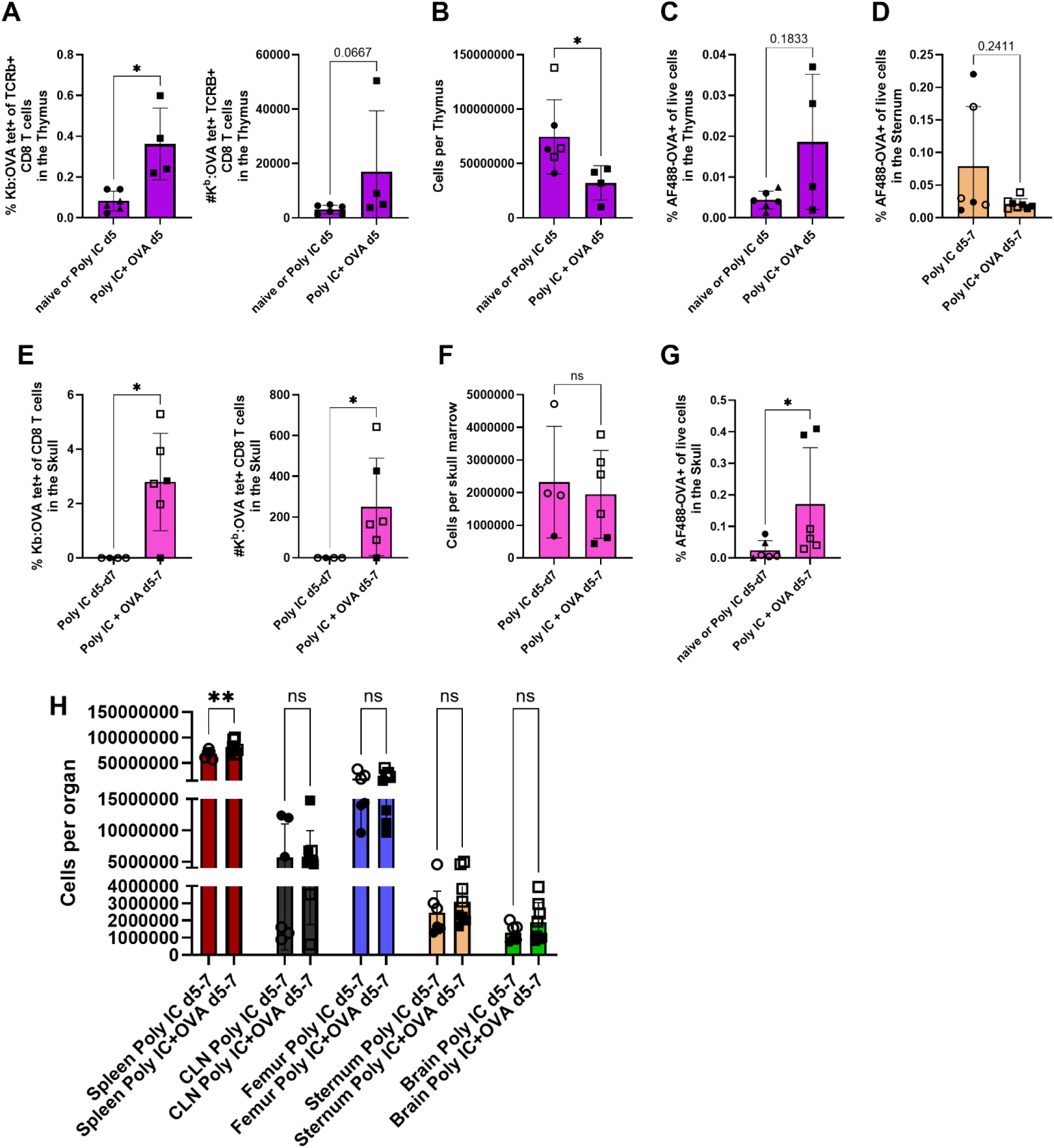
Acute inflammatory brain insults lead to the generation of antigen-specific CD8 T cells within the primary immune organs. Mice were injected with Poly I:C or Poly I:C+AF488-labeled OVA according to **Fig 3A**. In these mice, numbers of antigen-specific CD8 T cells and AF488+ cells in the thymus and bone marrow were evaluated. **A)** Frequencies and numbers of OVA-specific CD8 T cells in the thymus are compared in Poly I:C+OVA vs. Poly I:C alone groups. **B)** Total cells per thymus are shown. Injection of an antigen admixed with Poly I:C induces neuroinflammation which is associated with systemic thymic involution. Thymus tissue was only acquired from mice 5 days post-injection. Experiments in **A-B** are performed once on day 5. **C)** Frequencies of AF488+ cells of total live cells in the thymus are shown. **D)** Frequencies of AF488+ cells of total live cells in the sternal bone marrow are shown. **E)** Frequencies and numbers of K^b^: OVA specific CD8 T cells in the skull bone marrow are compared between Poly I:C+OVA and Poly I:C alone groups. Skull marrow has significant numbers of antigen-specific cells compared to controls. **F)** Total cells recovered from the skull bone marrow is comparable between groups. **G)** Frequencies of AF488+ cells of total live cells in the skull bone marrow are shown and found to be increased above control groups. Naïve mouse skulls were pooled in with Poly I:C control samples. Triangles represent naïve mice. **H)** Total cell counts (cells recovered from each organ) are shown. Organs were processed and immediately counted with trypan blue exclusion on a hemocytometer. These results are shown in **H**. All experiments in **D-H** were performed twice (once harvesting organs on day 5 and once on day 7 post injections into the brain). Data from both experiments are pooled. Open squares/circles are data from day 7, while closed squares/circles are data from day 5 post injury. Data are shown as individual mice. Mean and SD are shown. For comparisons in **A-G**, either a Welch’s t-test of a Mann Whitney test was used comparing the groups depending on normal or not-normal distribution of the data determined by the Shapiro-Wilk’s test. Comparisons made are shown on the graph. In **H**, a one-way ANOVA was performed the results of which were significant (P <0.0001). Then post-hoc analysis was performed using Šídák’s multiple comparisons test on the selected pairs shown on the graph. ns = P > 0.05, * = P ≤ 0.05, ** = P ≤ 0.01, *** = P ≤ 0.001, and **** = P ≤ 0.0001.

**Figure S4:**
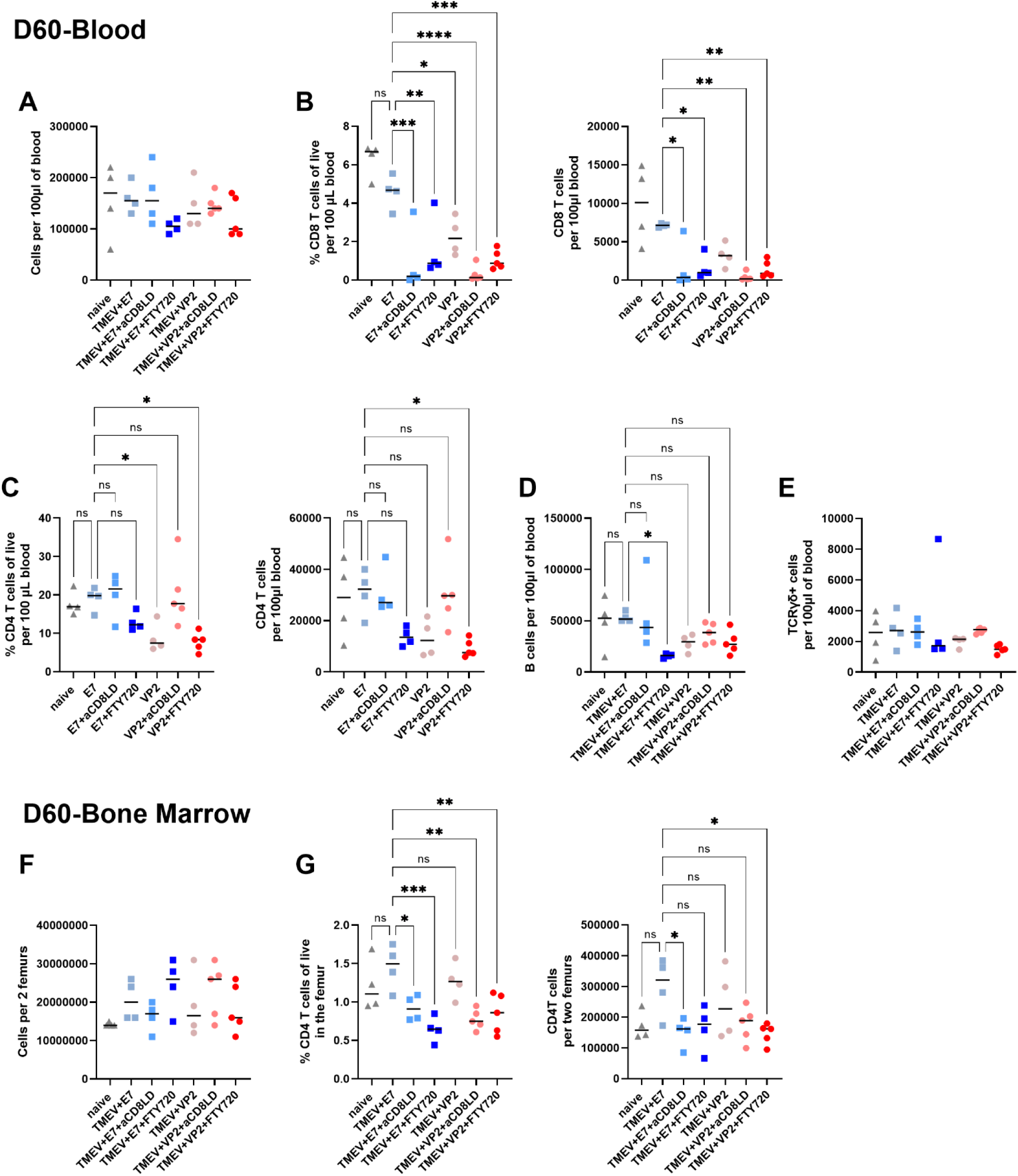
Virus antigen-specific CD8 T cells against neurotropic TMEV infection durably remain within the bone marrow tissue long-term. Experimentral design is as shown in **Fig 5A**. **A)** Total numbers of cells per 100 µl of blood is comaprable between groups. **B)** Differences in frequencies and counts of CD8 T cells in the blood are shown and found to be reduced in depleted and FTY720 treated groups, as expected. **C)** Minor differences in the frequencies and numbers of CD4 T cells between groups are observed. **D)** Total B cell counts are shown. **E)** No change in total counts of γδ T cells is seen in the blood 60 days post brain infection. **F)** Total counts of recovered cells per two femurs are comparable between groups. **G)** Frequencies and counts of CD4 T cells in the bone marrow across various groups are compared. **A-G** a one-way ANOVA followed by either Šídák’s multiple comparisons test or Dunnett’s multiple comparisons test was performed comparing between selected groups. If ANOVA results were not significant, no post-hoc analysis was performed. Data are shown as individual mice with mean and SD. Comparisons made are shown on the graph. ns = P > 0.05, * = P ≤ 0.05, ** = P ≤ 0.01, *** = P ≤ 0.001, and **** = P ≤ 0.0001.

**Figure S5:**
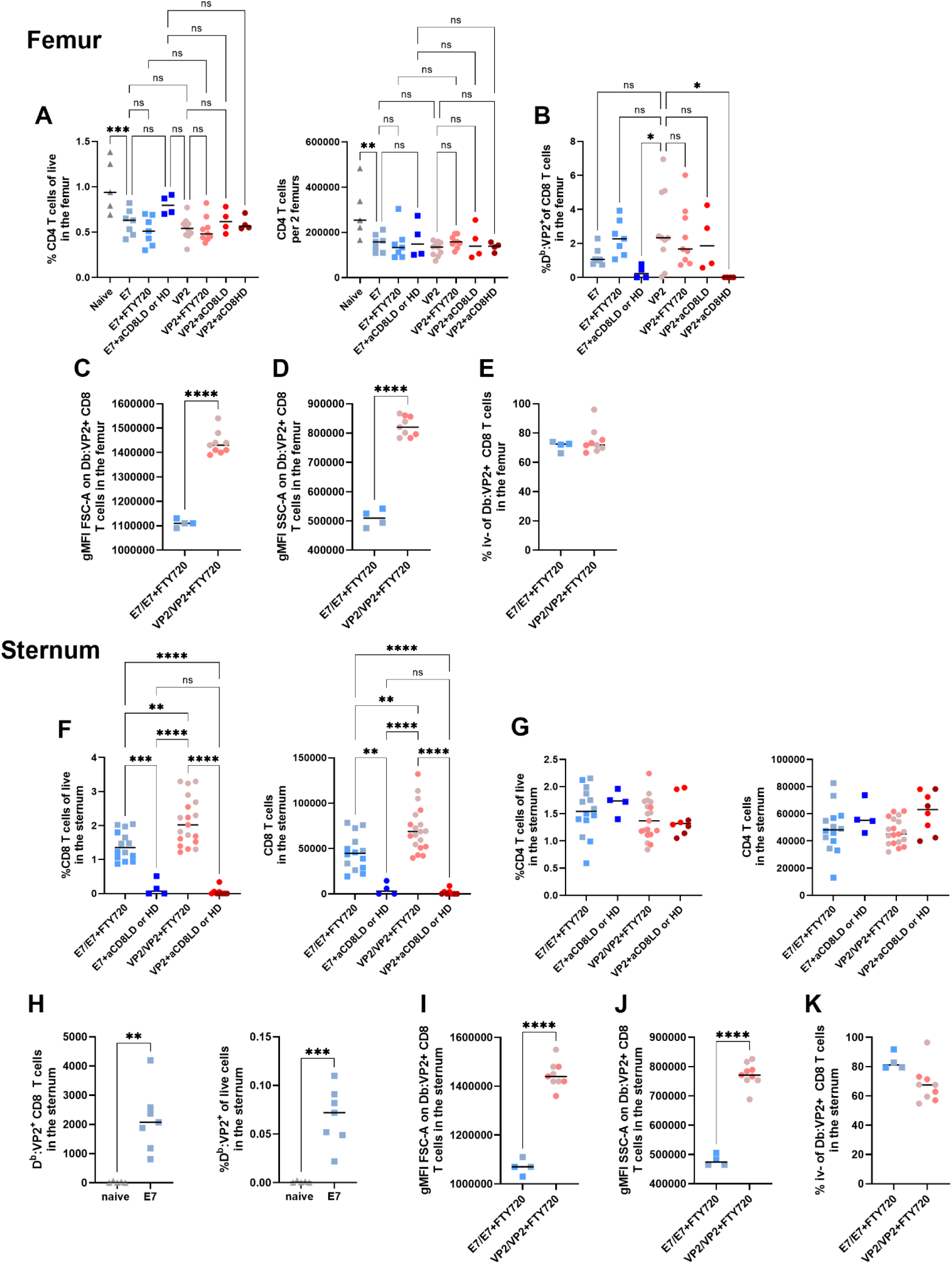
Virus antigen-specific CD8 T cells in the bone marrow can be reactivated upon antigen reencounter. Experimental design is as shown in **Fig 6A**. **A)** Frequencies and numbers of CD4 T cells in the femur are shown. **B)** Frequencies of antigen-specific CD8 T cells in the femur is compared amongst groups. **C-D)** Antigen-specific reactivation of memory CD8 T cells in the femoral bone marrow are associated with increased blasting as measured by gMFI of FSC-A and SSC-A. **E)** The majority of antigen-specific CD8 T cells within the femoral marrow are iv- indicating tissue residency. **F)** Frequencies and numbers of CD8 T cells in the sternal bone marrow are shown amongst groups. Pooled groups had no significant difference between them and retain previous color scheme from individual comparisons. **G)** No differences between groups are seen in frequencies or counts of CD4 T cells in the sternal bone marrow. **H)** At late memory time points, sternal bone marrow contains significant frequencies and numbers of viral antigen-specific CD8 T cells in quiescent previously infected mice compared to naïve never infected mice. **I-J)** Antigen-specific reactivation of memory CD8 T cells in the femoral bone marrow is associated with increased blasting as measured by gMFI of FSC-A and SSC-A. This is an independent experiment from data shown in **Fig 6I-J**. Experiments were performed once at 15 weeks and once at 28 weeks post infection and data are either pooled and shown or independently shown if gMFI is being compared. **K)** The majority of antigen-specific CD8 T cells within the sternal marrow are iv- indicating tissue residency. When comparing more than 2 groups, a one-way ANOVA with Šídák’s multiple comparisons test or Tukey’s multiple comparisons test was performed comparing between selected groups. Comparisons are shown on the graph. When comparing two groups, a Welch’s t test was performed. Color scheme in pooled groups (**C-G, I-K**) follows individual groups (**A-B**). Data are shown as individual mice with mean and SD. Comparisons made are shown on the graph. ns = P > 0.05, * = P ≤ 0.05, ** = P ≤ 0.01, *** = P ≤ 0.001, and **** = P ≤ 0.0001.

**Figure S6:**
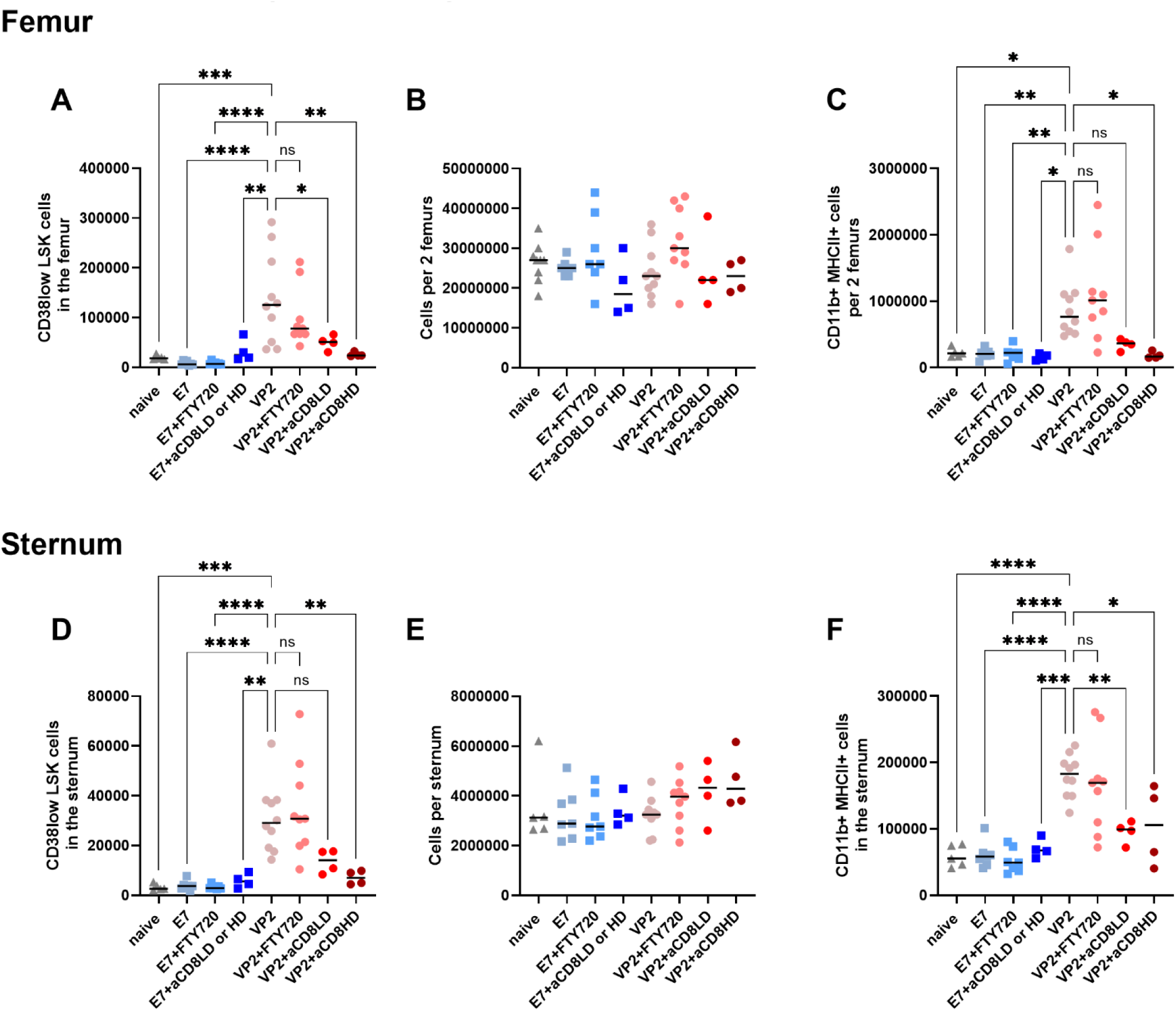
Hematopoietic stem and progenitor cells respond to the reactivation of antigen-specific memory CD8 T cells within the bone marrow. Experimental design is as previously described in **Fig 6A**. The stem cell populations from the same mice analyzed in Figure 6 and Figure 7 are shown here. **A)** Absolute counts of CD38 low LSK cells are increased following antigen-specific memory CD8 T cell reactivation in a CD8 T cell dependent manner in the femoral bone marrow. **B)** Total numbers of recovered cells per two femurs are comparable across groups. **C)** Absolute counts of CD11b+ MHCII+ cells in the femoral bone marrow are shown and indicate a CD8 T cell mediated effect upon antigen-specific memory CD8 T cell reactivation. **D-F)** similar results to the above are also seen in the sternal bone marrow. **D)** Absolute counts of CD38 low LSK cells are compared between groups. **E)** Comparable cell recovery per sternum is shown. **F)** Absolute counts of the CD11b+ MHCII+ cells in the sternal bone marrow are quantified. This influx in the sternum is abrogated when CD8 T cells are depleted. In **A-F**, a one-way ANOVA with Dunnett’s multiple comparisons test was performed comparing between selected groups. Comparisons made are shown on the graph. Data are shown as individual mice with mean and SD. If the results of ANOVA were not significant, no post-hoc analysis was performed. ns = P > 0.05, * = P ≤ 0.05, ** = P ≤ 0.01, *** = P ≤ 0.001, and **** = P ≤ 0.0001.

**Supplementary table 1:** Antibodies and specific reagents used.

| **Product** | **Dilution** | **Cat#** | **Company** |
| --- | --- | --- | --- |
| BUV395 αF4/80 | 1:200 | 565614 | BD |
| Zombie UV Viability Dye | 1:1000 | 423108 | Biolegend |
| BUV563 αSca-1 | 1:100 | 749200 | BD |
| BUV615 αNK1.1 | 1:100 | 751111 | BD |
| BUV737 αCD21-35 | 1:500 | 612810 | BD |
| BUV805 αMHCII (IA/IE) | 1:200 | 748844 | BD |
| BV421 αCD103 | 1:100 | 121422 | Biolegend |
| Pac Blue αCX3CR1 | 1:100 | 149037; 149038 | Biolegend |
| BV480 αCCR2 | 1:500 | 747971 | Biolegend |
| BV510 αCD4 | 1:50 | 100559 | Biolegend |
| BV570 αCD8α | 1:50 | 100740 | Biolegend |
| BV605 αCD11c | 1:100 | 117333; 117334 | Biolegend |
| BV650 αCD44 | 1:200 | 740455 | BD |
| BV711 αLy6G | 1:200 | 127643 | Biolegend |
| BV750 αMHCI | 1:200 | 746869 | BD |
| BV750 αckit/CD117 | 1:100 | 747412 | BD |
| BV786 αckit/CD117 | 1:100 | 564012 | BD |
| BV786 αMHCI | 1:500 | 749706 | BD |
| FITC αCD45 | 6 μg/<br>mouse<br>for iv<br>labeling | 35-0451-U100 | Tonbo (when we purchased<br>it was Tonbo, currently,<br>Cytex) |
| PerCP αLy6C | 1:100 | 128027 | Biolegend |
| BB700 αCD24 | 1:200 | 746122 | BD |
| PE αCD122 | 1:100 | 123210 | Biolegend |
| PE-CF594 αCD45 | 1:1000 | 562420 | BD |
| PE-Cy5 αCD11b | 1:1000 | 550112U100 | Tonbo |
| PE-Fire700 αCD38 | 1:100 | 102747; 102748 | Biolegend |
| PE-Cy7 αTCRβ | 1:100 | 605961u100 | Tonbo |
| SparkNIR 685 αCD69 | 1:200 | 104558 | Biolegend |
| AF700 αCD25 | 1:100 | 102024 | Biolegend |
| APC D<sup>b</sup>:VP2<sub>121-130</sub> tetramer | 1:100 | N/A | NIH tetramer facility |
| PE K<sup>b</sup>:SIINFEKL tetramer | 1:50 | TB-5001-1 | MBL life sciences |
| PE-H2Kb bound to<br>SIINFEKL | 1:50 | 141604 | Biolegend |
| APC-Fire 750 αCD62L | 1:100 | 104450 | Biolegend |
| APC-Fire 810 αB220 | 1:100 | 103277; 103278 | Biolegend |
| APC-Fire 810 αCD19 | 1:100 | 115578 | Biolegend |
| FC Block | 1:100 | 553141 | BD |
| BUV563 αCD69 | 1:100 | 741234 | BD |
| BUV661 αSca1 | 1:500 | 741466 | BD |
| BUV737 αCD103 | 1:100 | 569678 | BD |
| BUV805 αCD48 | 1:100 | 741945 | BD |
| Pacific Blue αCD62L | 1:50 | 104424 | Biolegend |
| BV480 αKit | 1:250 | 566074 | BD |
| BV510 αCD4 (this concentration used only in the OVA experiments) | 1:100 | 100559 | Biolegend |
| BV570 αCD11c | 1:250 | 117331 | Biolegend |
| BV605 αCD44 | 1:250 | 103047 | Biolegend |
| BV711 αCD45 | 1:250 | 103147 | Biolegend |
| BV750 αLy6G | 1:250 | 747072 | BD |
| RB613 αTCRβ | 1:250 | 758429 | BD |
| PerCP αLy6C (this concentration used only in the OVA experiments) | 1:250 | 128028 | Biolegend |
| BB700 αCD19 | 1:250 | 566411 | BD |
| PE-Cy5 αCD150 | 1:500 | 115912 | Biolegend |
| cFluor BYG750 αMHCII | 1:100 | R7-20563 | Cytex |
| PE-Cy7 αCD8 | 1:1000 | 100722 | Biolegend |
| APC αB220 | 1:250 | 103212 | Biolegend |
| Spark NIR 685 αNK1.1 | 1:500 | 156529 | Biolegend |
| APC-Cy7 αTCR$\gamma\delta$ | 1:100 | 118144 | Biolegend |
| APC-Fire 810 αCD11b | 1:500 | 101288 | Biolegend |
| **Other reagents** | **N/A** | **Cat#** | **Company** |
| Surface stain fixative. BD Cytofix | **N/A** | 554655 | BD |
| Percoll | **N/A** | P1644-1L | Sigma |
| FTY720 | **N/A** | SML0700-25MG | Sigma |
| Trypan Blue | **N/A** | 15250-061 | ROENZ |
| Ultracomp beads | **N/A** | 01-2222-42 | Invitrogen |
| Cytex QC beads (to ensure cytometer performance) | **N/A** | B7-10001 | Cytex |
| RPMI | **N/A** | 10040CV | Corning |
| RPMI | **N/A** | 11875093 | Gibco |
| PBS | **N/A** | 10010023 | Gibco |
| 10x PBS | **N/A** | 70011044 | GIBCO |
| Poly I:C | **N/A** | NC9180242 | Invivogen TLRLPIC |
| AF488-conjugated OVA | **N/A** | O34781 | Invitrogen |
| Sutures for wound closure post intracranial surgery for OVA injection experiments | **N/A** | ETHJ494G | Ethicon |
| Heparin sodium salt | **N/A** | H3393-50KU | Millipore/Sigma |

