## Supplementary figures and images for "Antigen-specific CD8 T cells are generated and reactivated in the bone marrow following viral brain infections and impact the bone marrow niche"

### Figure S1

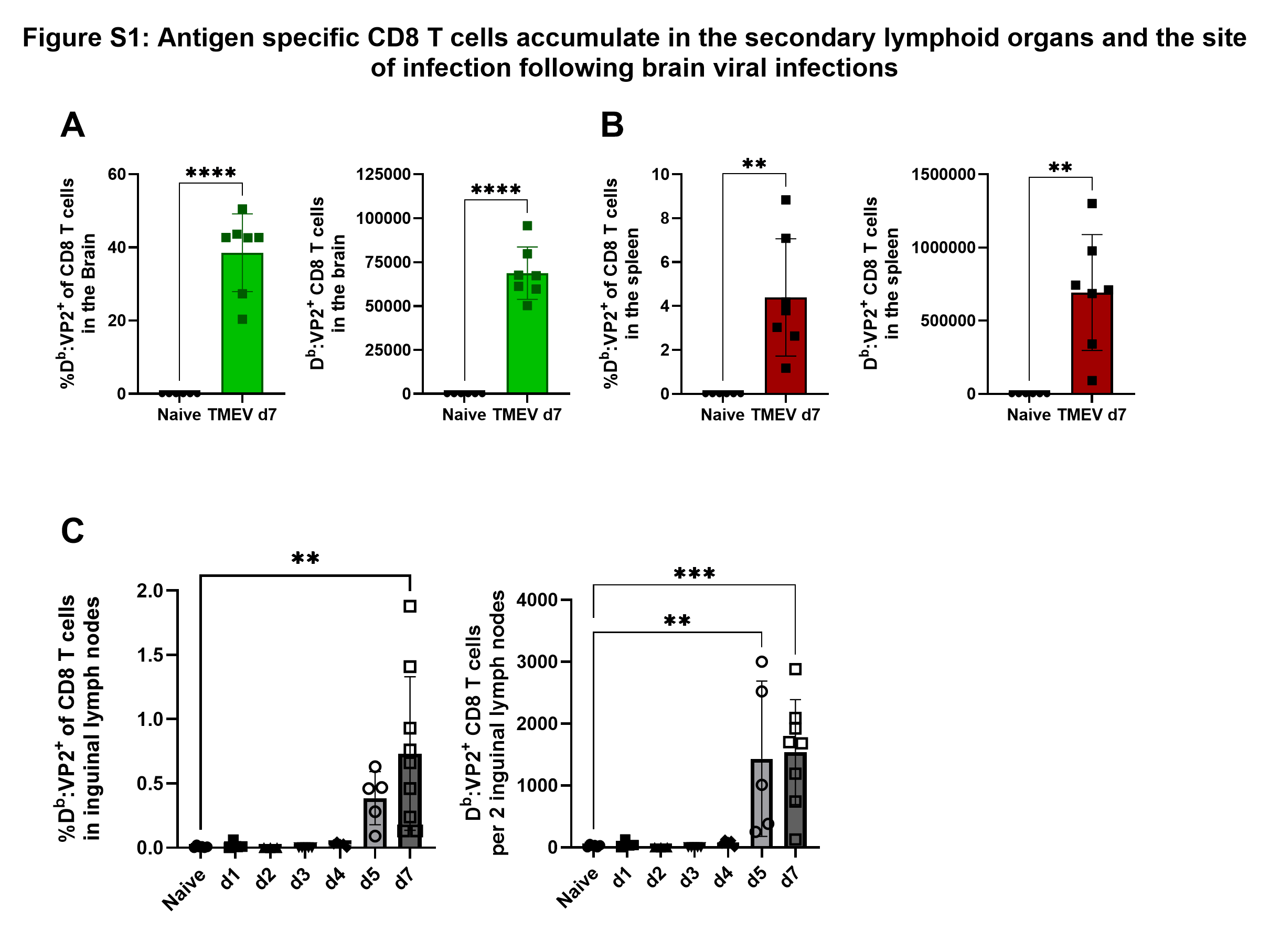

### Figure S2

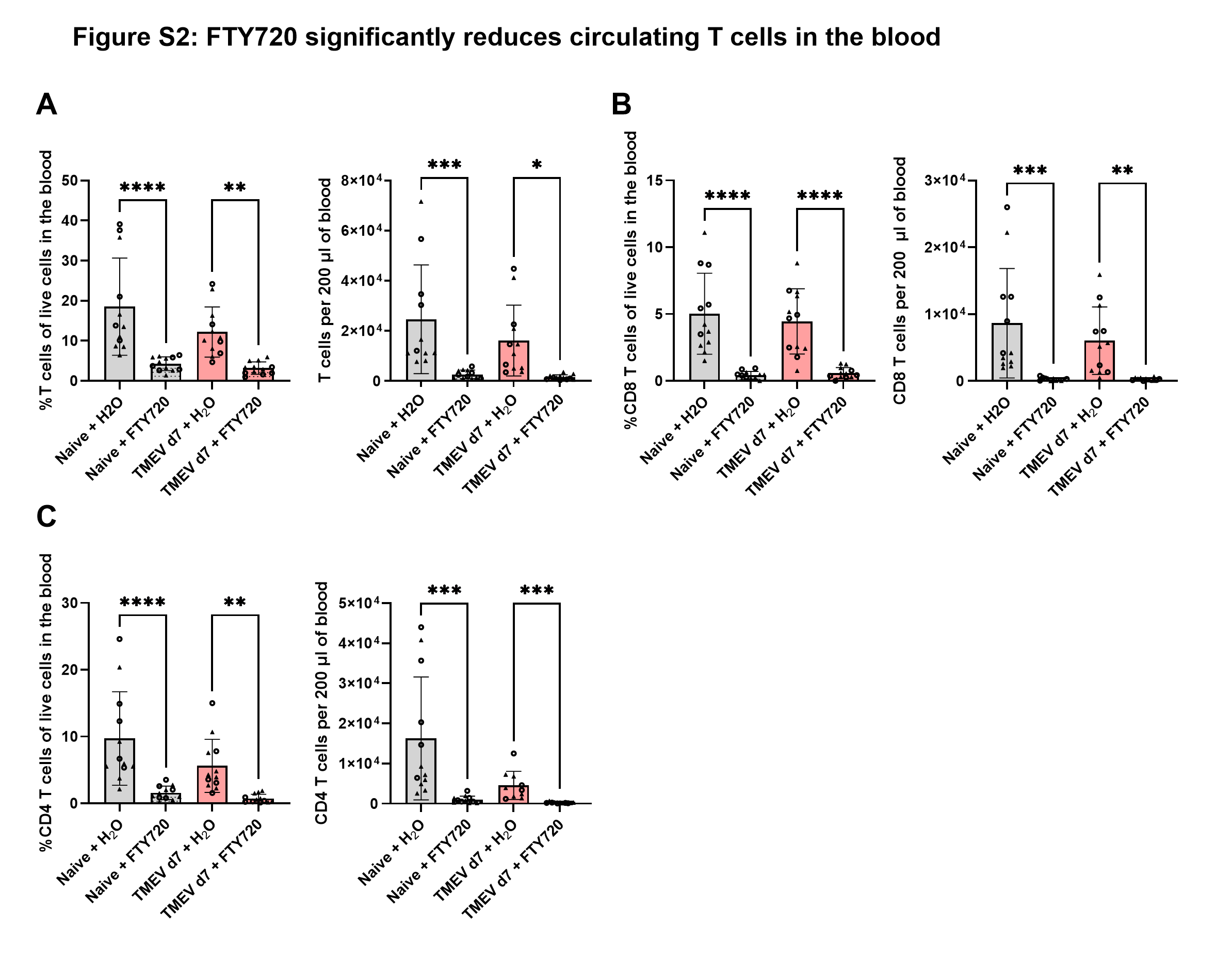

### Figure S3

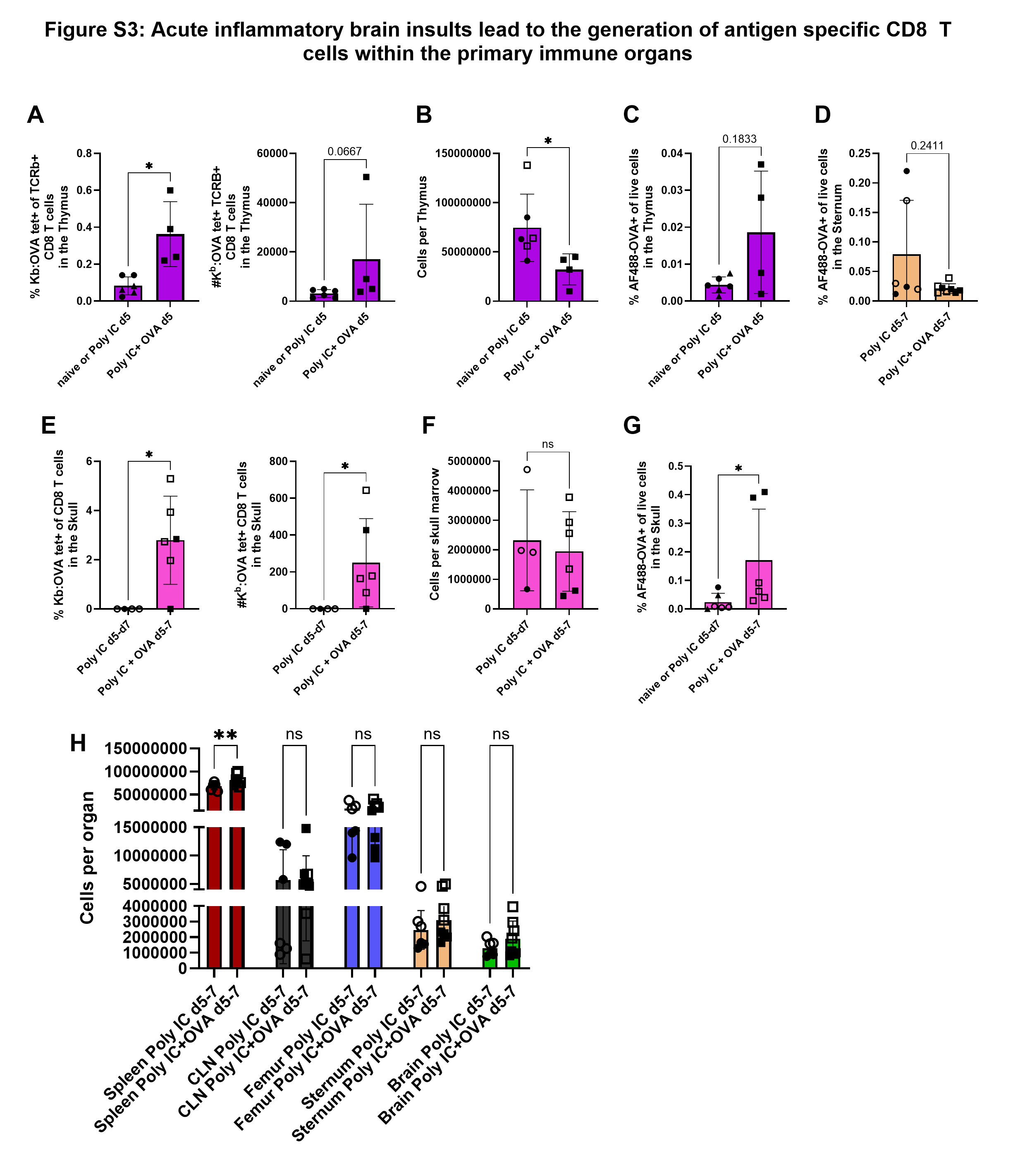

### Figure S4

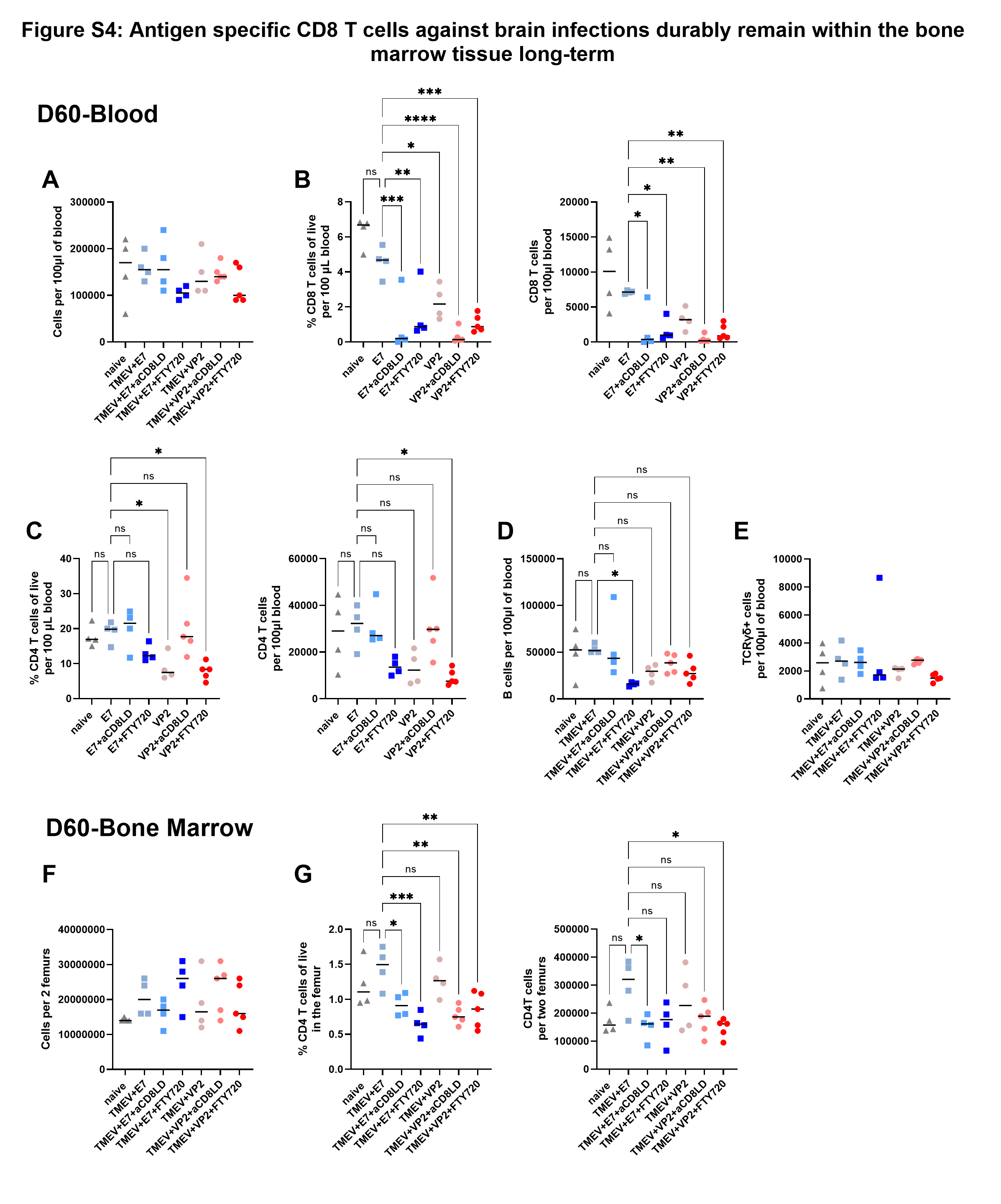

### Figure S5

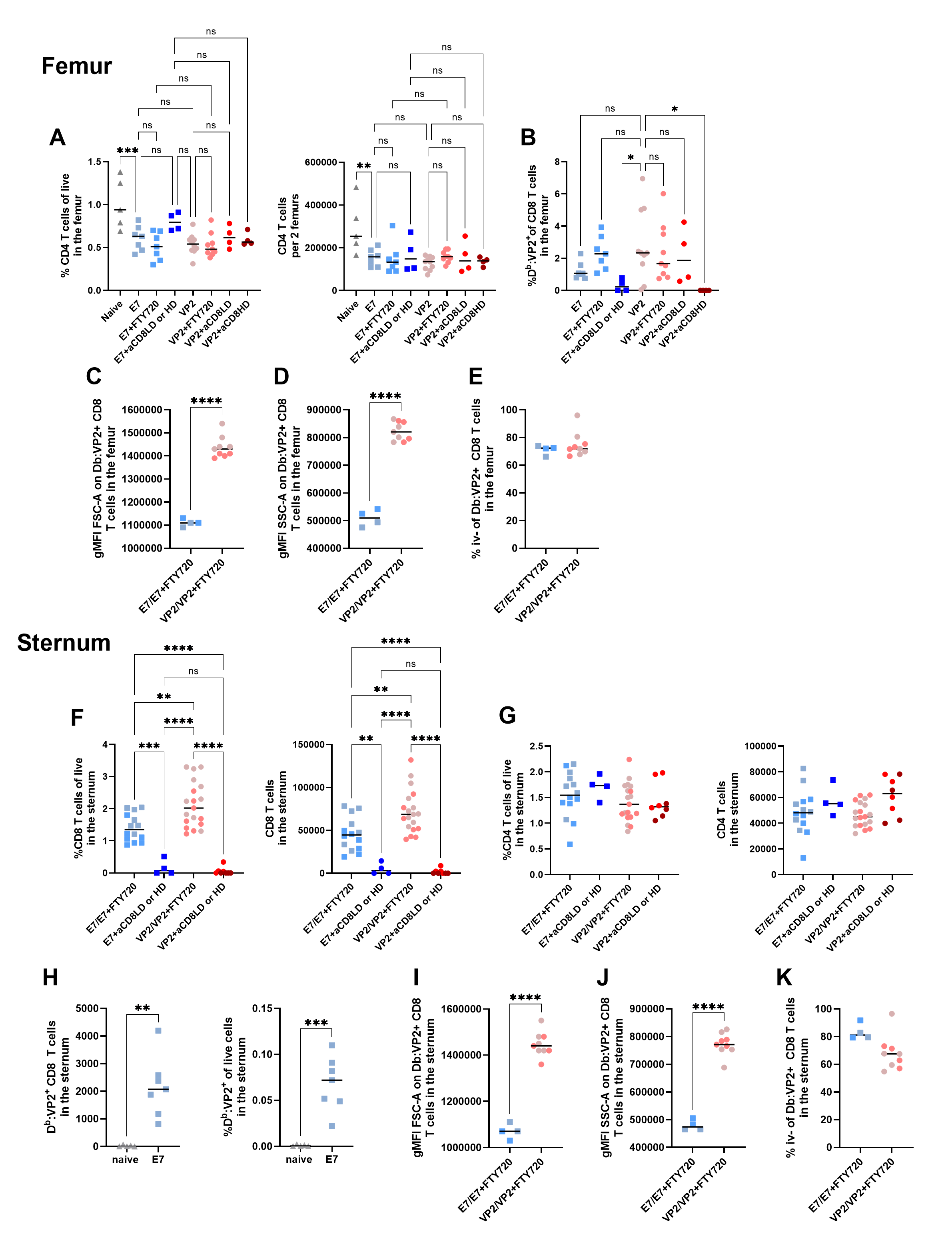

### Figure S6

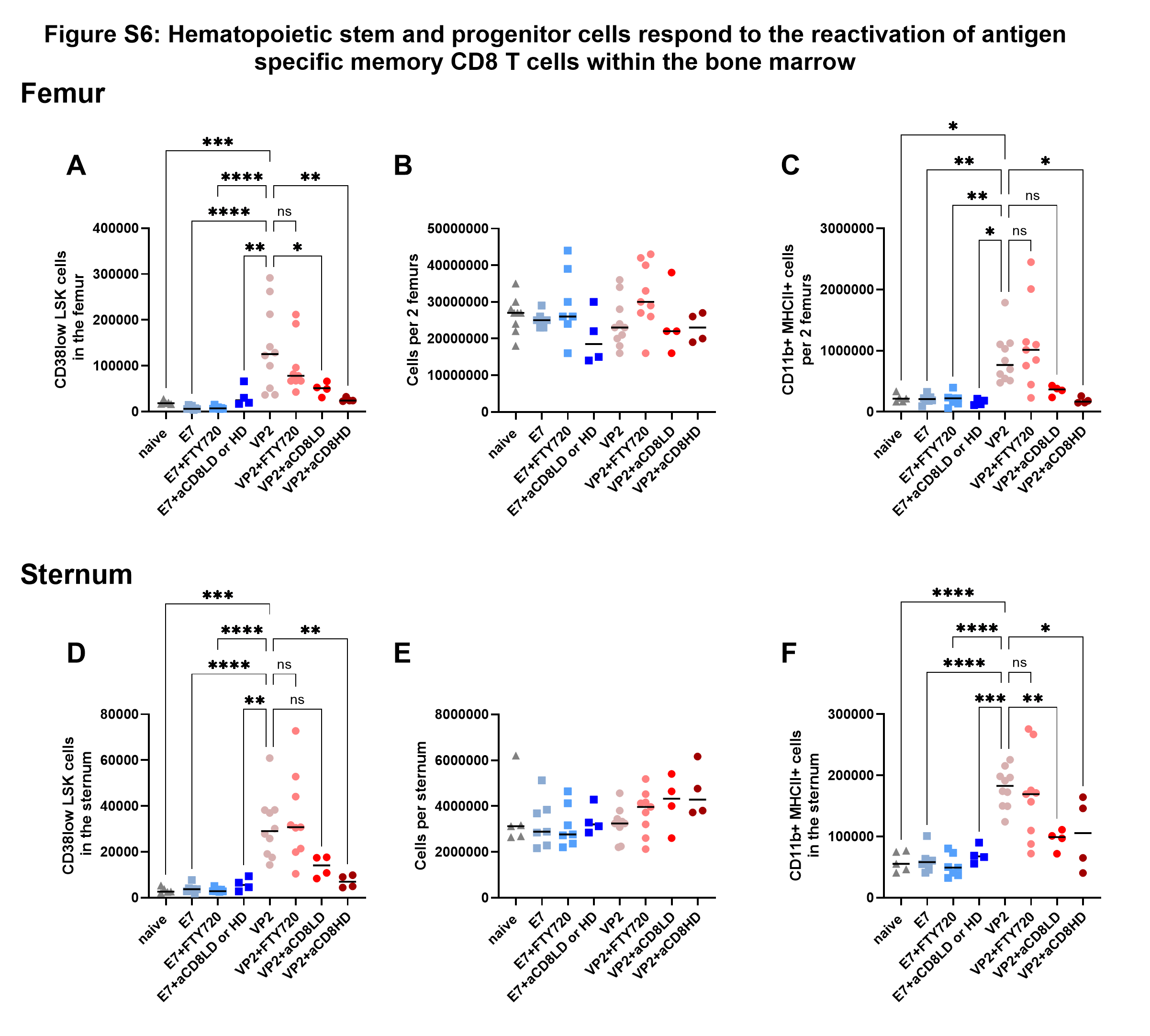
